# Insights into I-motif stabilization by high resolution primer extension assays: Its strengths and limitations

**DOI:** 10.1101/2022.01.05.475026

**Authors:** Jan Jamroskovic, Marco Deiana, Nasim Sabouri

## Abstract

Cytosine-rich DNA can fold into four-stranded intercalated structures, i-motif (iM), in acidic pH and require hemi-protonated C:C^+^ base pairs to form. However, its formation and stability rely on many other factors that are not yet fully understood. In here, we combined biochemical and biophysical approaches to determine the factors influencing iM stability in a wide range of experimental conditions. By using high resolution primer extension assays, circular dichroism and absorption spectroscopies, we demonstrate that the stability of three different biologically relevant iMs are not dependent on molecular crowding agents. Instead, some of the crowding agents affected overall DNA synthesis. We also tested a range of small molecules to determine their effect on iM stabilization at physiological temperature, and demonstrated that the G-quadruplex-specific molecule, CX-5461, is also a promising candidate for selective iM stabilization. This work provides important insights into the requirements needed for different assays to accurately study iM stabilization, which will serve as important tools for understanding iMs’ biological roles and their potential as therapeutic targets.

## Introduction

An important characteristic of the DNA molecule is its ability to form not only double-stranded DNA structures but also non-canonical DNA structures. Among these structures, cytosine-rich i-motif DNA structures (iM) were first identified *in vitro* in 1993 (Gehring et al., 1993), and many years later their formation in human nuclei was visualized by an iM-specific antibody (iMab) (King et al., 2020; Zeraati et al., 2018). Sequences that form iMs are for instance enriched in promoters of oncogenes such as BCL2, KRAS and MYC, and iM formations in these sites are suggested to act as molecular switches to control gene expression and to be modulated by small molecules (Kaiser et al., 2017; Kang et al., 2014; Kendrick et al., 2014; Sutherland et al., 2016).

iMs are formed in C-rich ssDNA by hydrogen bonding between hemi-protonated cytosines (C:C^+^). The pK_a_ of N3 position of the cytosine is ∼ 4.5 (Mergny et al., 1995), but other factors as characteristic DNA sequence and the length and number of cytosine tracts influence the formation and stability of iM in higher pH (Gurung et al., 2015; Wright et al., 2017). Also, chemicals and physical environmental factors have strong influence on iM formation. The most important factor is so far the pH of the environment, in which iM formation is heavily dependent on. Only slight pH changes affect iM folding and stability (Cheng et al., 2021a; Cheng et al., 2021b; Mergny et al., 1995). Other factors, such as temperature, ionic strength and crowding conditions have also positive/negative impact on the stability of the iM. Higher temperature and ionic strength cause iM destabilization, while crowding conditions favour their formation and increase stability (Dzatko et al., 2018; Mergny et al., 1995; Nguyen et al., 2017; Rajendran et al., 2010; Takahashi et al., 2017). Thus, mimicking the complex cellular milieu *in vitro* helps to accurately study iM, but all aforementioned factors must be strictly controlled for.

To characterize an iM structure and its folding, methods like ultraviolet (UV) absorption, nuclear magnetic resonance (NMR), and circular dichroism (CD) spectroscopies are widely used to record the typical iM spectral signature. The important parameter called pH_T_ (transition pH) is usually used to characterise iMs. It is defined as pH in which 50% of iM is folded at given temperature. Therefore, different iMs have different pH_T_ values that are determined experimentally by for instance absorption or CD measurements.

To reconstitute DNA replication *in vitro*, primer extension assay is commonly used. This experimental system consists of several components: i/ DNA template containing iM DNA sequence of interest with annealed fluorescently/radioactively labelled primer, ii/ DNA polymerase used to synthesize over the iM structure, iii/ dNTPs as DNA building blocks, iv/ Mg^2+^ cofactor and Na^+^ or K^+^ salts, and v/ BSA protein to avoid unspecific binding of DNA polymerase. Other additives as molecular crowders, or small molecules can be supplemented and their effect on iMs stability can be studied via DNA synthesis. The principle of the assay is to follow the extension of a labelled primer during DNA synthesis when annealed to a template DNA that can form different structures. Any disturbance/blockage during the synthesis will cause pausing/arresting of the DNA polymerase and eventually resulting in falling off the template and thus formation of shorter products than the full-length products. These products are separated on a denaturing polyacrylamide gel and later visualized (Scheme 1). Primer extension assays to study the stability of iMs was earlier reported by the Sugimoto lab. They showed that a fragment of the *E. coli* DNA polymerase II (Klenow fragment) pauses while replicating DNA containing iM-forming sequence (Takahashi et al., 2017). Various molecular crowders and co-solutes were used to mimic crowding effects of the cell inner environment, and thus achieve comparable conditions *in vitro.* pK_a_ of iMs was reported to be shifted under crowding conditions into neutral pH=7.0 and above. Another primer extension study was performed by the Muniyappa group where it was found that the DNA polymerase is paused at an iM-forming sequence from *S. cerevisiae* when treated by single-walled carbon nanotubes (SWNT), suggesting iMs pKa shift and stabilization in pH=7.5 (Kshirsagar et al., 2017).

In here, we employed high-resolution primer extension assay to step-wise monitor DNA synthesis in iM-containing templates. The products from the primer extension assay were separated on high resolution sequencing gels, which allowed us to follow DNA synthesis in one nucleotide steps over time in various conditions. When combined with proper controls, high-resolution primer extension assays are very informative, and the large sized sequencing gels allow running more than 50 samples in one gel which is advantageous when testing a number of various conditions, drugs and time points. For these assays, we selected some of the most stable iMs, which are reported to fold in neutral/basic pH and we tested their pH stability upon addition of co-solutes and molecular crowders (glycerol, PEG1000, PEG8000, dextran40 or ficol70), and upon treatment with various G4-stabilizing compounds. Furthermore, we have validated the primer extension assays with classical spectroscopic methods (absorption, CD, and NMR). Because iMs are highly dependent on the slight changes of pH, we paid special attention to carefully measure pH in all experiments, repeatedly. Every small change of pH was corrected and re-measured, especially when using different additives in high concentrations. Surprisingly, we did not detect any pK_a_ shifts into neutral pH in any of the used conditions and iM templates regardless of methods used (absorption, CD, NMRs, or primer extension assay). Moreover, we did not identify any iM destabilization in the primer extension assay that are caused by G4-ligands, instead we identified compound CX-5461 as a very potent iM stabilizer.

## Results and discussion

### Rapid unfolding of iM structures in pH titration

The iM-forming sequences, Jazf1, Dap and Hif-1α are present in the promoter regions of the *JAZF1* (JAZF Zinc Finger 1), *DAP* (Death-associated Protein 1) and *HIF1-α* (Hypoxia Inducible Factor 1A) genes. Jazf1 protein is a key transcriptional regulator of ribosome biogenesis and insulin translation, and it is associated to prostate cancer and endometrial stromal tumors (Kobiita et al., 2020; Koontz et al., 2001; Thomas et al., 2008). Dap and Hif-1α are also associated to cancer and correlate with metastasis of breast cancer (Ebright et al., 2020; Wazir et al., 2015). All three iM-forming sequences were earlier reported to be stable at neutral and basic pH, (Dzatko et al., 2018; Takahashi et al., 2017; Wright et al., 2017). However, a more recent study conducted in the Mergny lab did not confirm previously published pH_T_ values and showed that Dap and Hif-1α are unstable at physiological conditions (pH 7.0) (Cheng et al., 2021a).

To investigate the reason of these discrepancies between the published reports and to determine the stability of these iMs, we first employed spectroscopic techniques (CD, UV absorption, and ^1^H NMR) to measure iM folding at 25 °C. It is known that iMs are highly dependent on the environmental pH and minor changes might have immediate effect on the structure formation, therefore we paid extra care for the selection of buffer, ionic strength (Mergny et al., 1995), and temperature in our measurements. We selected MES buffer with pKa at 6.21 because the buffer range is suited for iM studies, and it is widely used in the field of non-B DNA research. Furthermore, throughout the whole study, the ionic strength was calculated and maintained with KCl at the constant level of 145 mM. To confirm the correct pH of each sample, pH was measured immediately before and after recorded CD and absorption spectra. Recorded CD spectra in pH=5.4 showed characteristic spectral signatures for iM formation with a positive peak signal at 288 nm and a negative peak at 255 nm for all three IM sequences (Figure S1). When increasing pH from 5.4 to higher pH, the CD peak at 288 nm decreased, and a clear transition between iM and non-iM DNA was observed through the characteristic formation of a well-defined isodichroic point centred at 276.5 nm. Similarly, the absorption band at 295 nm decreased with increasing pH (Figure S1). Changes of CD_max_ (CD peak at 276 - 288 nm) and Abs_295_ were plotted against pH to determine the pH_T_ values (Figure 1 and S2). Unfolding curves clearly showed two slopes, suggesting heterogenous formation of different iM structures in different pH. The slope was less steep in lower pH, but after a critical value the slope became steeper, and the structure unfolded quickly. Therefore, two pH_T_ values were calculated by fitting the data into bi-dose response function (Figure S2). For simplification, also one overall pH_T_ was determined by fitting the data into a single dose-response function (Figure 1). The 1D ^1^H NMR spectra for all three sequences showed the characteristic iM signals around 15-16 ppm at pH=6 or lower, which disappeared at pH =7.2 (Figure 1C), confirming our observed analysis by CD and UV absorption (Figure 1A-C). In summary, all three sequences folded into iMs, however, none of them were stable at pH=7.0 or higher, which is in agreement with studies conducted by the Mergny lab (Cheng et al., 2021a).

**Figure 1.**
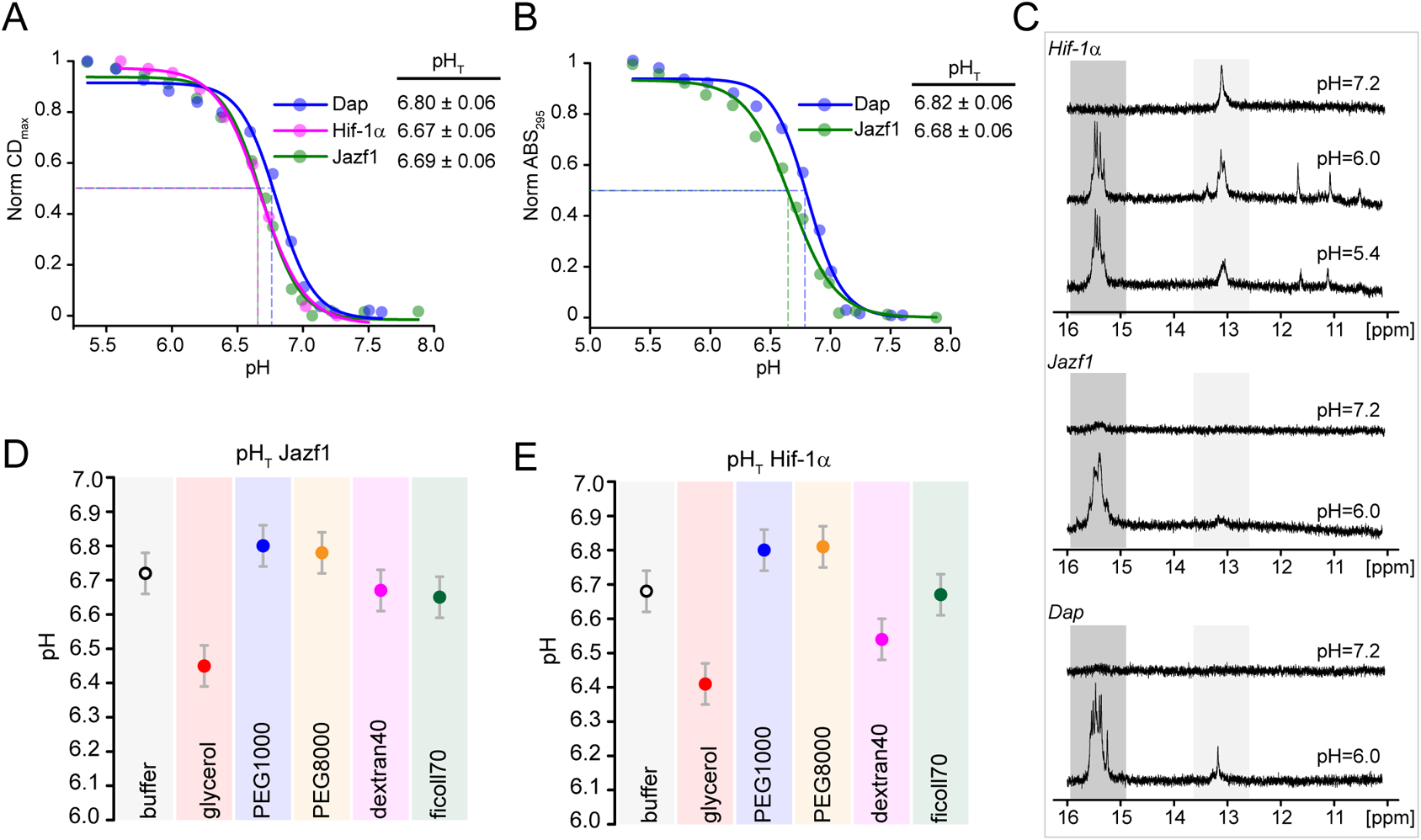
IMs fold at close-to-neutral pH at 25°C. Fitting curves of CDmax (A) and ABS295 (B) of iMs spectra recorded at 25 °C. Dotted lines in A and B show a pH_T_ transition point for each curve. (C) Identification of structural species formed within DNA used in the study. Imino proton region of the 1D ^1^H NMR spectra of free DNA in Hif1α (top panel), Jazf1 (middle panel) and Dap (low panel) sequences, pH of each sample is depicted above each spectrum, spectra were recorded at 25 °C, concentration of DNA was 170 μM. Dark grey panel shows region for iM, light grey panel show region for Watson-Crick base pairing. Graphical representation of pH_T_ values of Jazf1 (D) and Hif-1α (E) iMs in crowding conditions calculated from CD_max_ values. Original spectra for A, B, D and E can be found in Figure S1.

Next, we examined iM formation during *in vitro* DNA replication in primer extension assays at 37 °C (Han et al., 1999b). For this assay, the different DNA templates containing iM-forming sequences were annealed to a fluorescently labelled DNA primer. The final templates consist of dsDNA part following by 10 nucleotide ssDNA linker to ensure no steric interference between the DNA polymerase and the folded iM (Scheme 1). Ionic strength was kept constant at 145 mM in all reaction conditions by addition of KCl, and the pH was measured by a microprobe to ensure correct pH of the reaction environment. For the primer extension assay, we used all three iM-sequences, Jazf1, Dap and Hif-1α, and a non-iM forming control sequence, which was derived from *S. pombe* genome. To be able to follow the whole process of DNA synthesis in all reactions with different pH, samples were collected over 40 minutes, and visualized on 10% denatured polyacrylamide sequencing gel which allows 1 nt resolution of the newly synthesized DNA.

Despite of the decreased polymerase activity in pH below 6 (Richard et al., 2006), the amount of pausing signal can still be attributed to the stability of the iM in various pH. We observed strong pausing right before an iM-forming sequence already after 0.5 min of the reaction in pH=5.6 with all three templates. For the Jazf1 template, two pausing regions were detected. The first region was at nucleotide 10, just before the start of the C-rich iM sequence. With progression of time, pausing was also detected at 11^th^ and 12^th^ position, suggesting that the JazF1 iM slows/pauses DNA synthesis, but that continuous reloading of the polymerase allows bypassing of the iM. The second pausing site was observed at the 3^rd^ C-tract, suggesting that another type of iM folding or an alternative type of structure may be present. The pausing sites were reduced during the time course of the reactions, however still some pausing was detected even after 40 minutes. With increasing pH and time of the reaction, the pausing sites were less pronounced, and finally no pausing sites were detected after 0.5 min in pH=6.4, resulting in maximum full-length products almost immediately after the start of the reaction (Figure 2 and S3). Similar observations were made with Dap and Hif-1α templates, but these iMs were fully bypassed without any detectable pausing already at pH=6.2. Similar additional pausing site as Jazf1 before the second C-tract was observed in the Dap template, but not in the Hif-1α template (Figure S4, S5). Control reactions with non-iM DNA template did not show specific pausing sites, but an overall slower DNA synthesis at pH=6 and lower (Figure S6), which agrees with previous reports (Richard et al., 2006). To quantify stability of iMs during DNA synthesis in primer extension assay, maximum pausing signal from individual reactions was plotted over the pH and pH_T_ values were calculated representing 50% iM stability (Figure 2E). PH_T_ values of Dap and Hif-1α were 6.0 and they were fully unfolded in pH=6.2 and pH_T_ for Jazf1 was 6.2 and fully unfolded in pH=6.4. The stability of the iM structures in the primer extension assays showed a similar steep slope as unfolding in CD and absorption pH titration experiments. Jazf1 showed the best stability in the given pH and had the best resolution for detection of pausing during the course of the primer extension reaction.

**Figure 2.**
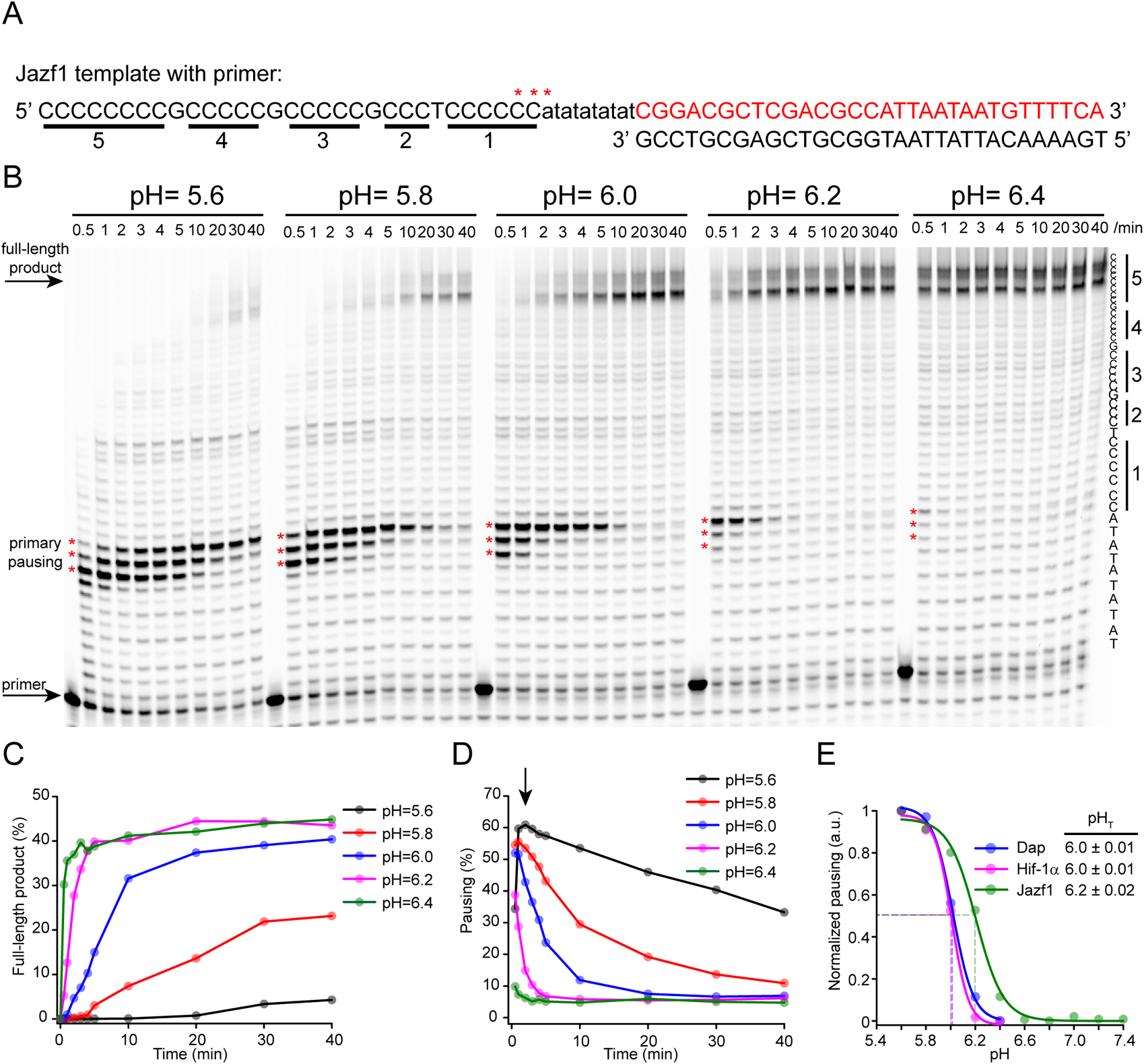
Jazf1 iM pauses DNA synthesis in high resolution primer extension assay. Sequence (A) and representative sequencing gel (B) of primer extension assay with template containing Jazf1 iM sequence, pH of the reaction is written above each gel. Time-points (min) when individual samples were collected is described above each well. C-tracts from iM sequence are underlined in A and depicted on the right side of the gel. Pausing sites are depicted as red stars (***) in each reaction and shown also in the sequence in A. Quantification of full-length product (C) and pausing signal (D) of each reaction is describing an effect of pH on overall reaction and iM, respectively. Normalized pausing of primer extension assay with all three iM templates as a function of pH (E).

### Molecular crowders and co-solutes do not shift the pK_a_ of iMs

It is reported that molecular crowders and co-solutes shift the pK_a_ of N3 in the cytosine base leading to increased stabilization of iMs in higher pH (Bhaysar-Jog et al., 2014; Rajendran et al., 2010; Takahashi et al., 2017). Polyethylene Glycol (PEG) are used as the main preference for those experiments, however a clear mechanism of how PEGs act on iMs remains unknown. Generalization of these phenomena led us to investigate the possible effects of co-solutes and crowders on the pK_a_ shift of Jazf1 and Hif-1α iMs. Therefore, we recorded CD spectra of iMs supplemented with 20% (v/v) of glycerol, PEG1000 or 20% (w/v) of PEG8000, dextran40 or ficoll70. The current pH of each sample was confirmed before and after each recorded CD spectrum. CD_max_ at 288 nm and transition to ssDNA and 276.5 nm was not affected by any of the tested additives (Figure S7 and S8). Next, we plotted CD_max_ over pH and calculated the pH_T_ values (Figure 1C, D). All pH_T_ values in the presence or absence of additives were in the range of 6.7 ± 2SD for both Jazf1 and Hif-1α iMs. Addition of glycerol affected pH_T_ slightly more, pH_T_ of Jazf1 and Hif-1α was 6.45 and 6.41, respectively. Overall, differences in pH_T_ values were very subtle, suggesting that neither of these agents shifts the pK_a_ of the tested iM sequences to neutral values.

### Molecular crowders and co-solutes do not cause selective replication fork blockage at the iM

Since we did not detect a shift of iMs pK_a_ to neutral pH in our CD measurements, we instead tested the hypothesis that iMs in crowding conditions are stabilized and hence enhance the pausing of the DNA polymerase before the iM. We selected Jazf1, the most stable iM, and performed primer extension assays at pH=6.0. The processivity of the polymerase at pH=6.0 was not strongly affected in this assay. Neither was primer consumption affected and no random pausing was observed in the control reaction (Figure S6). We performed reactions supplemented with 20% (v/v) of glycerol, PEG1000 or 20% (w/v) of PEG8000, dextran40 or ficoll70. Some of the additives affected pH, therefore the composition (basic and acidic part of the buffer) of each reaction was recalculated, and pH was adjusted to pH=6.00 ± 0.05. Ionic strength was again kept constant at 145 mM by addition of KCl (Figure 3 and Figure S9). The pausing sites in Jazf1 template on the position 10, 11 and 12 detected in the buffer only control after 0.5 min, was also detected in the presence of glycerol, dextran40, and ficoll70. When reaction progressed, pausing at 12^th^ nucleotide remained visible during the first five minutes, but the iM was fully bypassed after 10 minutes. PEG1000 caused pausing in the linker region as well and the primer was not extended enough to reach the iM in the first 0.5 min. Therefore, pausing was detected at later time points. In these reactions, the primers were fully consumed only after three minutes of reaction. PEG8000 also inhibited the start of the reaction, and most of the primer was not utilized by the polymerase even after 40 min of reaction (Figure S9). On the other hand, CD peaks at 288 nm were not affected by the different PEGs or the other additives (Figure 1D). Therefore, slower replication by PEGs suggest potential inhibition through the binding of the polymerase to the primer-template, instead of selective iM stabilization. Alternatively, it decreases the processivity of the polymerase or precipitate the polymerase. Furthermore, only 20% maximum pausing signal was detected in the PEG1000 sample compared to 45% ± 5 in all the other reactions (Figure 3G).

**Figure 3.**
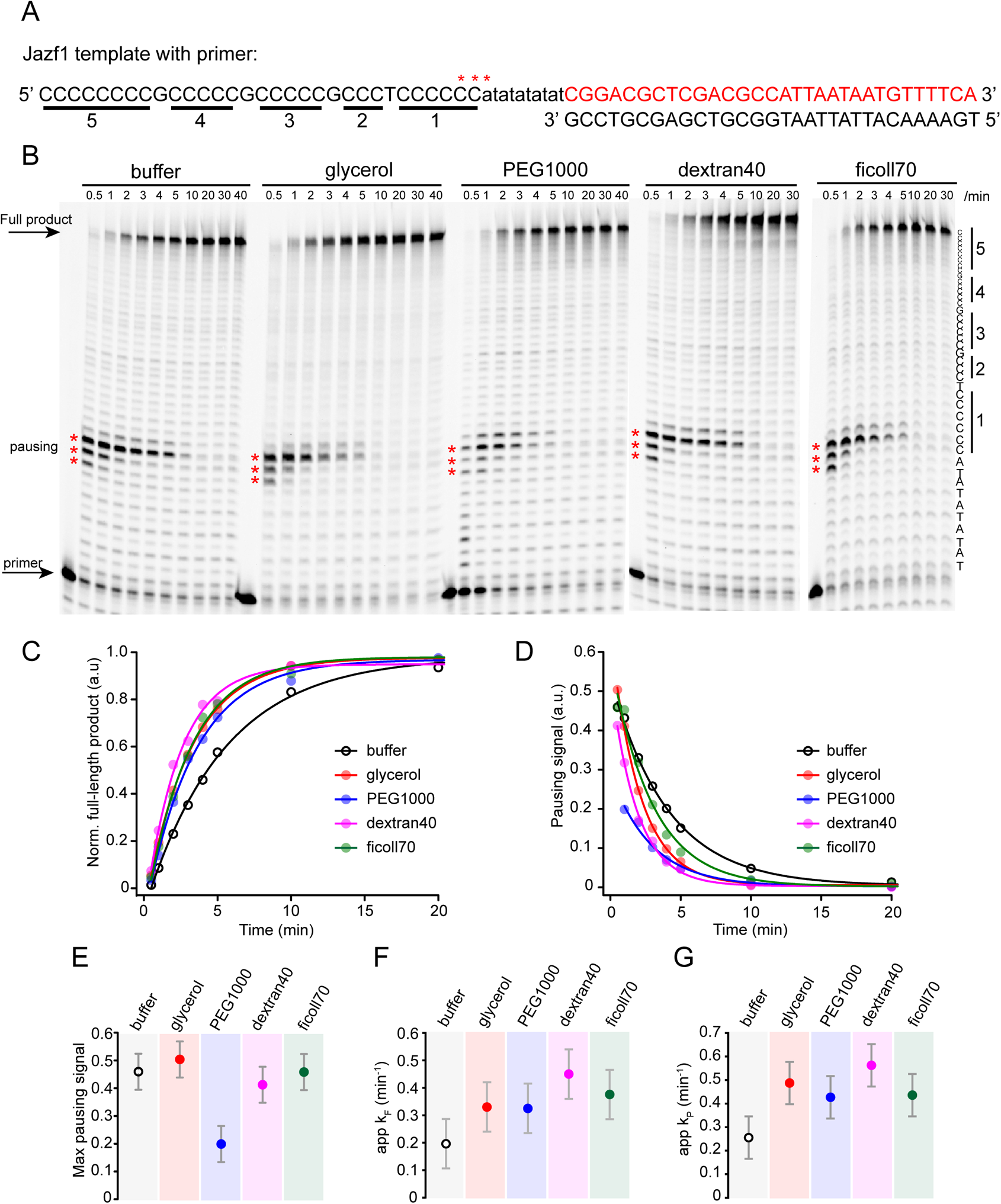
Molecular crowders and co-solutes do not increase the stability of Jazf1 iM structure in high-resolution primer extension assay, but facilitate the reaction. Sequence (A) and representative sequencing gel (B) of primer extension assay with template containing Jazf1 iM sequence. Reactions were supplemented with co-solutes or molecular crowders, specified above each reaction. Time of sample collection (min) is written above each well. C-tracts from iM sequence are underlined in A and depicted on the right side of the gel. Pausing sites are depicted as red stars (***) in each reaction and are also marked in A. Quantified full-length products of all reactions were normalized to the maximum signal (C). Pausing signals of individual reactions were subtracted by their minimal signal (D). Proportion of maximum pausing signal for each condition was compared (E). Data from individual reactions were fitted into exponential equation of the first order in OriginLab software and apparent rate constants for synthesis of full-length product k_F_ (F) and rate for disappearance of pausing site k_P_ (G) were calculated and. All data are shown as mean ± pooled SD.

Both the synthesis of the full-length product and the decreased pausing signal were plotted over time, and exponential equations were used to mathematically fit the data. Apparent rate constants for a particular reaction were derived for the synthesis of full-length product (k_F_) and for the decay of a pausing signal (k_p_). We compared k_F_ and k_P_ among all the tested conditions (Figure 3E, F). All additives increased both k_F_ and k_P_ values compared to a control reaction suggesting faster progression (processivity) of the polymerase, but neither stabilizing nor destabilizing effect of the Jazf1 iM was observed. These results were also observed with the Dap and Hif1-α templates (Figure S11 and S12).

Reaction supplemented with PEG1000 had a peculiar behaviour. The primer was utilized slower, the pausing sites were detected later in the reaction and the maximum pausing signal was also decreased, but k_F_ and k_P_ were higher than in the control with no additives. As shown above, CD spectra from pH titrations were not affected by PEG1000. So, this behaviour is caused by affecting the binding of the DNA polymerase to DNA or its processivity. This effect was even more pronounced by PEG8000.

### CX-5461 stabilizes iMs in primer extension assay

Earlier reports have shown that some G4 ligands also bind iMs, and even cause iM destabilization (Abdelhamid et al., 2019; Pagano et al., 2018). Therefore, we examined the stabilization/destabilization behavior of a few ligands in our high-resolution primer extension assay. We anticipated that compounds which destabilize iM will facilitate DNA synthesis over iM by decreasing polymerase pausing, on the other hand compounds stabilizing iM will enhance polymerase pausing before the iM. We tested various concentrations of G4 ligands including: TmPyP4 (Han et al., 1999a), PhenDC3 (Chung et al., 2014), pyridostatin (Rodriguez et al., 2008), CX-5461 (Xu et al., 2017), CX-3543 (Drygin et al., 2009), Braco19 (Read et al., 2001), and quinazoline 8g (Jamroskovic et al., 2020). We also used the iM stabilizer, ellipticine-based NSC71795 (King et al., 2020) as a positive control and the DNA intercalator, BMH-21 (Peltonen et al., 2010) as a negative control. The concentration range used in the experiments was optimized for each compound. We selected the most stable iM Jazf1 and ran primer extension reaction in pH=6 for four minutes. After four minutes, pausing sites at the iM is still clearly visible (20-30% of fluorescence signal) (Figure 2B-gel pH=6), and allows us to follow both stabilization and destabilization effects of the iM by the ligands. None of the ligands facilitated DNA replication passed the iM by destabilizing the structure, as the pausing sites in all reactions had similar intensity. However, in high concentrations many of the compounds affected replication in an unspecific manner by interfering with DNA synthesis. This included, NSC71795 that unspecifically affected both, iM and non-iM templates (Figure S12 and S13). Only two compounds CX-5461 and Braco19 caused increased pausing of the DNA polymerase close to the iM structure position at pH=6. Moreover, CX-5461 did not affect replication over non-iM DNA even at 50 μM concentration, but Braco19 affected iM and Non-iM DNA replication at 25 μM. We selected both these compounds for more in detail examination and tested their stabilizing potential in various pHs with three different templates, Jazf1, Hif-1α and non-iM (Figure 4). Here, we decreased the time of reaction to 1 minute to be able to follow maximum polymerase pausing, which occurred in the beginning of the reaction. After one minute, we observed pausing at nucleotides 10, 11 and 12 on the Jazf1 template (Figure 2B-gel pH=6) and at nucleotides 14 and 15 on the Hif-1α template (Figure S5). In DMSO-treated reaction, these pausing sites disappeared with increasing pH, firstly at nucleotides 10 and 11 of Jazf1 at pH=6. At pH=6.4, pausing was not detected. Similarly, both pausing sites at nucleotides 14 and 15 of Hif1α template disappeared quickly with increasing pH. CX-5461 enhanced the pausing of the DNA polymerase on nucleotides 10, 11 and 12 of Jazf1 template until pH=6.2. Similarly, CX-5461 stabilized Hif-1α iM in pH=5.8 and 6.0, but pausing was dramatically decreased at pH=6.2.

**Figure 4.**
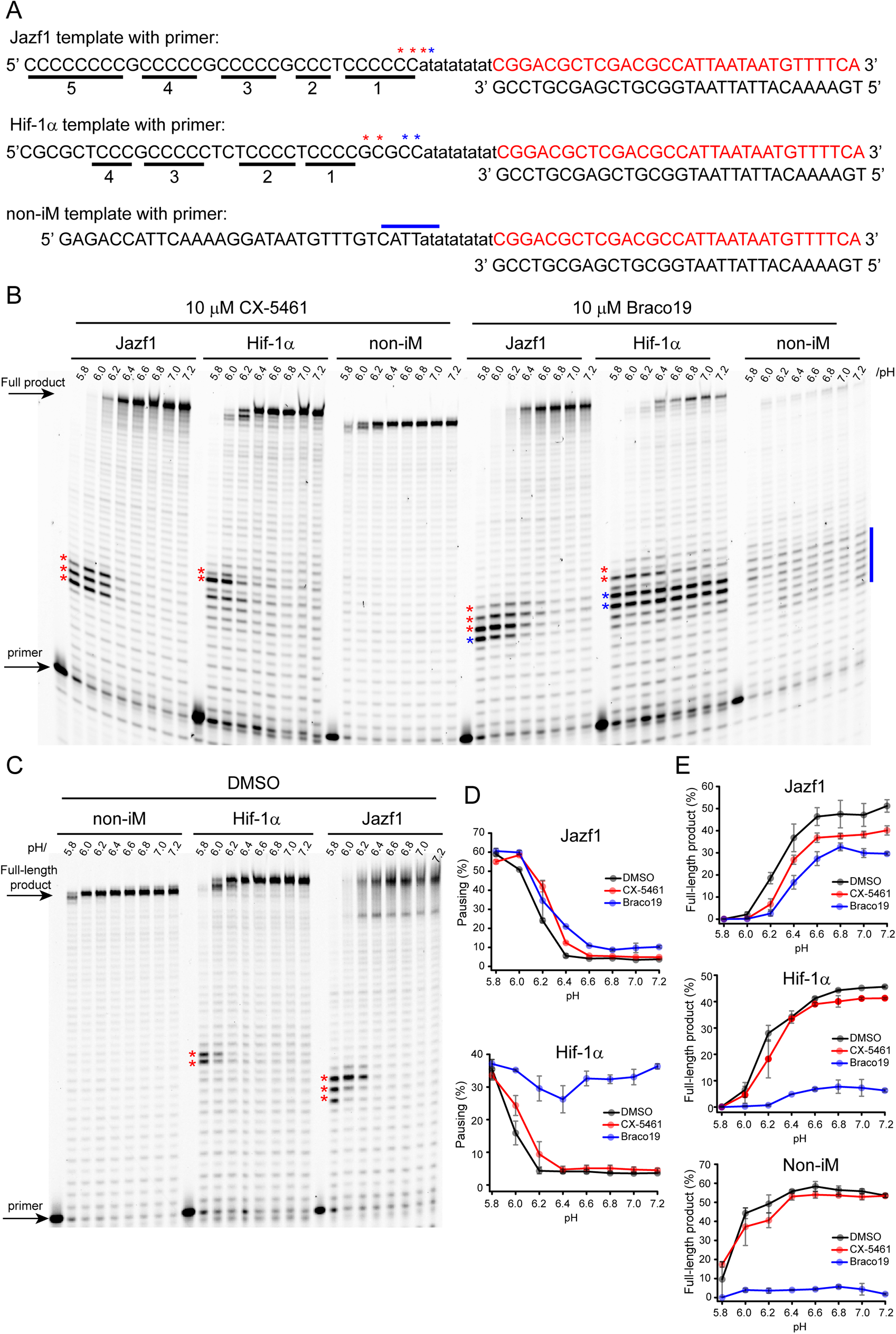
CX-5461 specifically stabilizes iMs in high resolution primer extension assay. Sequence (A) and representative sequencing gels (B, C) of primer extension assay with template containing Jazf1 iM sequence. Pausing sites are depicted as red star in each gel in B and C and in DNA sequence in A. Additional pausing sites formed upon binding of Braco19 are depicted as blue star in the gels and in DNA sequence in A. Blue line in non-iM DNA sequence in A and gel in B shows polymerase pausing caused by Braco19. IM containing template is written above each gel set and pH of each reaction is depicted above each well in B and C. Quantification of pausing signal (D) represents the sum of the signal depicted by red stars for DMSO and CX-5461 reaction and blue stars in case of Braco19 treatment. Quantification of full-length product (E) for all three DNA templates.

For the Braco19-treated reactions, a new strong pausing site at nucleotide 9 in the Jazf1 template was detected and pausing was still visible at pH=6.4. In the Hif-1α template, pausing at nucleotides 11 and 12 was very strong and persisted throughout all pHs until pH=7.2. But replication of the non-iM template was also strongly affected, with little amount of full-length products due to several pausing sites between nucleotides 8-14. This effect on the non-iM template was not visible after the four minutes reaction (Figure S13) due to continues recycling by the DNA polymerase that was allowed during the longer reaction time. Formation of additional pausing sites in the presence of Braco19 on Hif-1α template suggests that Braco19 binds other types of non-iM structures and causes polymerase pausing, and therefore can be attributed to a false positive result, and not to inducing iM stabilization. ^1^H NMR spectra of Hif-1α DNA showed two peaks, one characteristic for iM at 15-16 ppm and second peak characteristic for Watson-Crick base pairing at 13 ppm (Figure 1C). The second peak at 13 ppm was present throughout all tested pHs, at pH 5.4, 6.0 and 7.2, while the iM peak was not present at pH=7.2 anymore. To determine if Braco19 binds to other types of structures, we recorded ^1^H NMR spectra upon it’s binding to Hif-1α DNA. At pH=5.4, the specific signal at 13 ppm disappeared when using equimolar concentrations of Braco-19, suggesting its interaction to the non-iM structure. Titration experiments with Braco19 were performed at pH=6.0 and 7.2. Increased concentration of Braco-19 resulted in diminishing signal at 13 ppm for both pH=6.0 and 7.2, supporting our hypothesis that Braco19 interacts with the non-iM structure in the Hif-1α template (Figure 5). Our conclusion is also supported by observed binding of Braco19 to dsDNA (Deiana et al., 2021).

**Figure 5.**
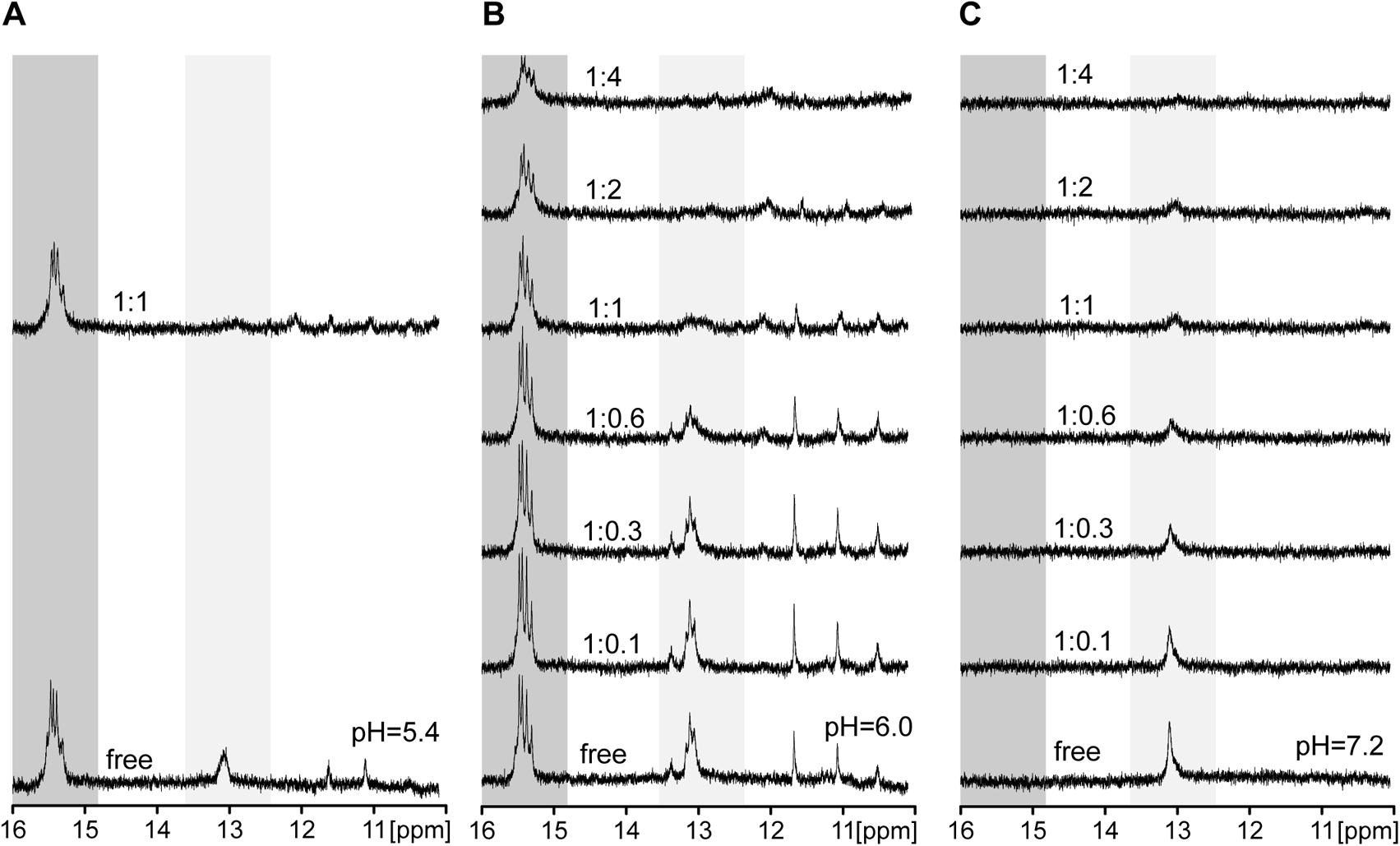
Braco19 interacts with both non-iM and iM structures folded within the Hif-1α DNA region. Imino proton regions of the 1D ^1^H NMR spectra of Hif-1α DNA in pH=5.4 (A), pH=6.0 (B) and pH=7.2 (C) titrated with Braco19. Spectra were recorded at 25 °C. Concentration of DNA was 170 μM. Ratio of DNA:Braco19 is indicated above each spectrum. Dark grey panel shows region for iM, light grey panel shows region for Hairpin structure.

## Discussion

High-resolution primer extension assay is a very valuable tool to study the effect of iMs on DNA replication. It’s one nucleotide resolution allows identification of specific pausing sites and how they resolve or are bypassed in time. Primer consumption, and synthesis of the full product give additional information about the reaction. Possible confounders can be identified when all necessary controls are made, including time-resolution.

Here, we have used primer extension assay to compare pH stability of three different iMs and their effect on DNA polymerase. We identified polymerase secondary pausing site in Jazf1 and Dap DNA templates, which suggests heterogeneity in iM folding. We re-assessed the effects of 20% molecular crowders on the stability of iMs in primer extension assay, which was suggested by Sugimoto (Takahashi et al., 2017), but we did not confirm any stabilizing effect when using 20% crowders. Rather, PEGs affected primer consumption, and reactions were inhibited in an iM-independent manner. It was also suggested that some of the typically used G4-ligands bind to iM and destabilize them (Abdelhamid et al., 2019; Pagano et al., 2018). In that case, we anticipated that DNA replication would be facilitated upon treatment with those compounds. But majority of the compounds had no effect on the stability of the iMs, but high concentrations unspecifically affected replication of both iM and non-iM templates, which is not uncommon for small molecules. Only one of the tested compounds CX-5461 specifically stabilized iMs and non-iM replication was not affected. We believe that further examination of CX-5461 and its analogues can be very useful to identify potent iM stabilizers.

C-rich DNA sequences containing self-complementary parts may preferably use Watson-Crick base pairing to fold to an alternative structure, usually hairpins (Kaiser et al., 2017; Kang et al., 2014; Kendrick et al., 2014; Sutherland et al., 2016). In our case Hif-1α sequence contain self-complementary regions on both sides, and we confirmed the existence of Watson-Crick base pairing by ^1^H NMR. This alternative structure of Hif-1α was present in neutral and basic pH and co-existed together with iM in lower pHs. We associated it with the polymerase pausing during Hif-1α replication, which caused false positive results upon Braco19 treatment in primer extension assay.

The limitation of the high-resolution primer extension assay is that it does not measure stability of the iMs directly, rather via activity of the polymerase during DNA replication. Therefore, it is not straightforward to identify possible confounders affecting other components of the reaction. Special attention must be paid to primer consumption and to the reaction kinetics. High concentration of some additives may inhibit the reaction start and pausing will appear later in time mimicking iM stabilization, which was the case with PEGs. Another limitation, while using high concentrations of molecular crowders, is increased viscosity of the samples making it problematic to manipulate with, to resuspend and to load into the polyacrylamide gel. High concentrations of some crowders, for instance dextran40 and ficoll70, even influence the migration of the DNA into the gel. Therefore, samples must be diluted 5-20x, which might affect the limit of detection of some of fluorescent dyes used to label the primer. In our case, Tet-label was sufficient. Furthermore, low volumes of the assay make it difficult to correctly measure pH of the samples, hence using special pH microprobe for small volume is very beneficial.

Finally, adopting high-solution primer extension assay, and coupling it with other classical spectroscopic methods, for instance CD, NMR, and absoprtion allowed us to study iM stabilization in detail. This combination serves as another important tool in understanding the biological roles of iMs and their folding patterns. Mimicking cellular milieu is especially valuable to make a correct connection between *in vivo* (Dzatko et al., 2018; King et al., 2020; Wright et al., 2017) and *in vitro* biophysical experiments (Cheng et al., 2021a; Cheng et al., 2021b) and can answer important fundamental biochemical questions.

## Material and methods

### Material

Lyophilized DNA molecules were purchased from Eurofins Genomics Ltd (Germany) and dissolved in deionized water according to manufacturer instructions. All DNA sequences used in the study are listed in the Supplementary information in Table S2.

50% (v/v) stock solutions of glycerol (MW = 92.09, MP Biomedicals, LLC), PEG1000 (MW ≈1000, Sigma), and 50% (w/v) of PEG8000 (MW ≈ 8000, Sigma), dextran40 (MW = 40 000, TCI) and Ficoll 70 (MW = 70 000, GE Healthcare) were prepared in deionized water and stored at 4 °C in dark to prevent their degradation.

### Measurement of pH

All pH measurements were performed by pH meter Mettler Toledo Five Easy Plus (Metler Toledo). Samples with bigger volumes for CD and absorption spectroscopy were measured by DG 115-SC probe (Mettler Toledo), temperature was manually set to 25 °C, all samples were incubated in 25 °C water bath (Julabo TW12). Samples with small volumes of 300 μL used for primer extension assays were measured by In Lab Micro-probe ISM (Mettler Toledo) equipped with temperature probe, incubated in 37 °C water bath (Julabo TW12) until temperature of the samples reached to at least 36 °C.

Both pH probes were calibrated by two-point calibration at pH=4 and pH=7 with freshly purchased calibration buffers (Mettler Toledo) prior to each measurement in the same temperature, volume, and container as was used to measure samples. Standard measurement error of pH meter referred by manufacturer was ± 0.005 pH units.

### CD and absorption measurements

For CD and absorption measurements, 1.4 μM DNA diluted in deionized water was heated to 95 °C for 10 minutes and slowly cooled down overnight to room temperature to melt all possible secondary structures coming from DNA lyophilization. Next day, DNA was diluted to final concentration of 0.5 μM into MES buffer. Composition of acidic and basic component of MES buffer of the specific pH at 25 °C was calculated using an online platform from Centre for Proteome Research, Liverpool (https://www.liverpool.ac.uk/pfg/Research/Tools/BuffferCalc/Buffer.html). Ionic strength was kept constant at 145 mM in all samples by KCl. pH of each sample was measured. Additives, glycerol and PEG1000 in final concentration of 20% (v/v) and PEG8000, dextran40 and ficoll70 in 0.2 mg/ml, were added into the sample, and pH of each sample was re-measured. If additives changed pH, then buffer composition was recalculated, and new samples were prepared with final pH required for measurement. Buffer composition was set as final when desired pH was within standard deviation of ± 0.05 pH units.

CD and absorption spectra were recorded at 25 °C by Jasco J-810 spectropolarimeter (Jasco International Co. Ltd.) using 1 cm path length quartz cuvette. Each spectrum was an accumulation of three scans recorded over 230-350 nm with an interval of 0.5 nm at a scan rate of 100 nm x min^−1^. Baseline for each measurement was corrected by subtracting the signal from sample without DNA. All spectra were smoothed in Origin Lab 2016 Software for visualisation purpose. Each sample in pH titration was prepared separately to control for ionic strength (145 mM). Ionic strength of additives was assumed to not to be a contributing factor to ionic strength.

### Primer extension assay

Tet-labelled primer (1 μM) was annealed to DNA templates (1.25 μM) in 50 mM KCl at 95 °C for 5 minutes in aluminium heating block and slowly cooled down overnight. Annealed DNA in final concentration of 40 nM was incubated in the buffer containing 50 mM MES in required pH, 8 mM MgCl2, 1 mg/ml BSA, 0.05 U of Klenow fragment (Thermo Scientific) and KCl in the final ionic strength of 145 mM in each sample depending on the specific pH. pH was measured multiple times in 300 uL volume and final pH was determined with SD of 0.05 pH units. Samples were incubated at 37 °C in water bath for 10 minutes to equilibrate, and reactions were started by addition of 0.2 mM dNTPs. Each reaction ran in 100 uL volume. Samples were taken from the main reaction in designated time-points (described in each well in the sequencing gels) and transferred into a STOP solution (95% formamide, 20 mM EDTA, and 0.1% bromophenol blue) in 1:1 ratio. A total of 6 uL was loaded on the sequencing gel containing 10% polyacrylamide, 8 M urea, 25% formamide, and 1× standard TBE buffer (Tris/borate/ethylenediaminetetraacetic acid). The gel was visualized with a Typhoon Scanner 9200 (GE Healthcare) at the Alexa setting of *λ* = 532 nm and quantified with the ImageQuant TL software (GE Healthcare). Primary pausing site was expressed as a sum of the total signal at nucleotides 10, 11 and 12 for Jazf11 template, 14 and 15 for Hif1-α and 10 and 11 for Dap template.

Samples containing co-solutes and molecular crowders: The final pH of each sample was measured to be 6.00 ± 0.05 after addition of co-solutes and molecular crowders: 20% (v/v) of glycerol, PEG1000, or 0.2 g/ml PEG8000, dextran40 and ficoll70. All primer extension reactions were run and sampled as described above.

Compound testing: All reactions were supplemented with various concentrations of small molecules: CX-3543, CX-5461, NSC71795 (ellipticine analogue), PhenDC3, pyridostatin (PDS), TmPyP4, Braco19, BMH-21 and quinazoline 8g (a kind gift from the Chorell lab, Umeå University). All concentrations are listed in the figures. 5% DMSO was used as control reaction. Reactions ran for four minutes at pH=6.0 in initial screening. For pH screening, reactions were run for one minute with 10 μM CX-5461, 10 μM Braco19 or 5% DMSO.

### Nuclear magnetic resonance measurements

1 mM iM stock solutions were prepared by adding deionized water into lyophilized DNA, heated to 95 °C for 5 min and slowly cooled down before an experiment. An effective DNA concentration of 170 μM was obtained by mixing DNA, 50 mM MES buffer (pH=5.4, 6.0 or 7.2 at 25 °C) and KCl into 154 mM ionic strength and 10% D_2_O. NMR samples were prepared in 3 mm NMR tubes. Samples were titrated by adding small volumes of 10 mM Braco19 solubilized in water, titration steps are described directly in the figure. All spectra were recorded at 298 K on a Bruker 850 MHz Avance III HD spectrometer equipped with a 5 mm TCI cryoprobe. Excitation sculpting was used, and 256 scans were recorded. Processing was performed in Topspin 4.1.3 (Bruker Biospin, Germany).

### Data fitting

For pH_T_ values, the middle point of the pH titration curves derived from CD and absorption measurements and from primer extension assays was obtained by fitting the data into dose-response function. In case of expressing two slopes, data were fitted into bi-dose response function in OriginLab 2016 software and pHT_1_ and pHT_2_ was determined.

Apparent rate constants of full product synthesis k_F_ and iM decay k_P_ were obtained by fitting the data into one-phase exponential decay (ExpDec) function in OriginLab 2016 software: *y* = *y*_0_ + *Ae*^−*x*/*t*^ from where apparent rate constant is defined as *k* = 1/*t*_1_.

---

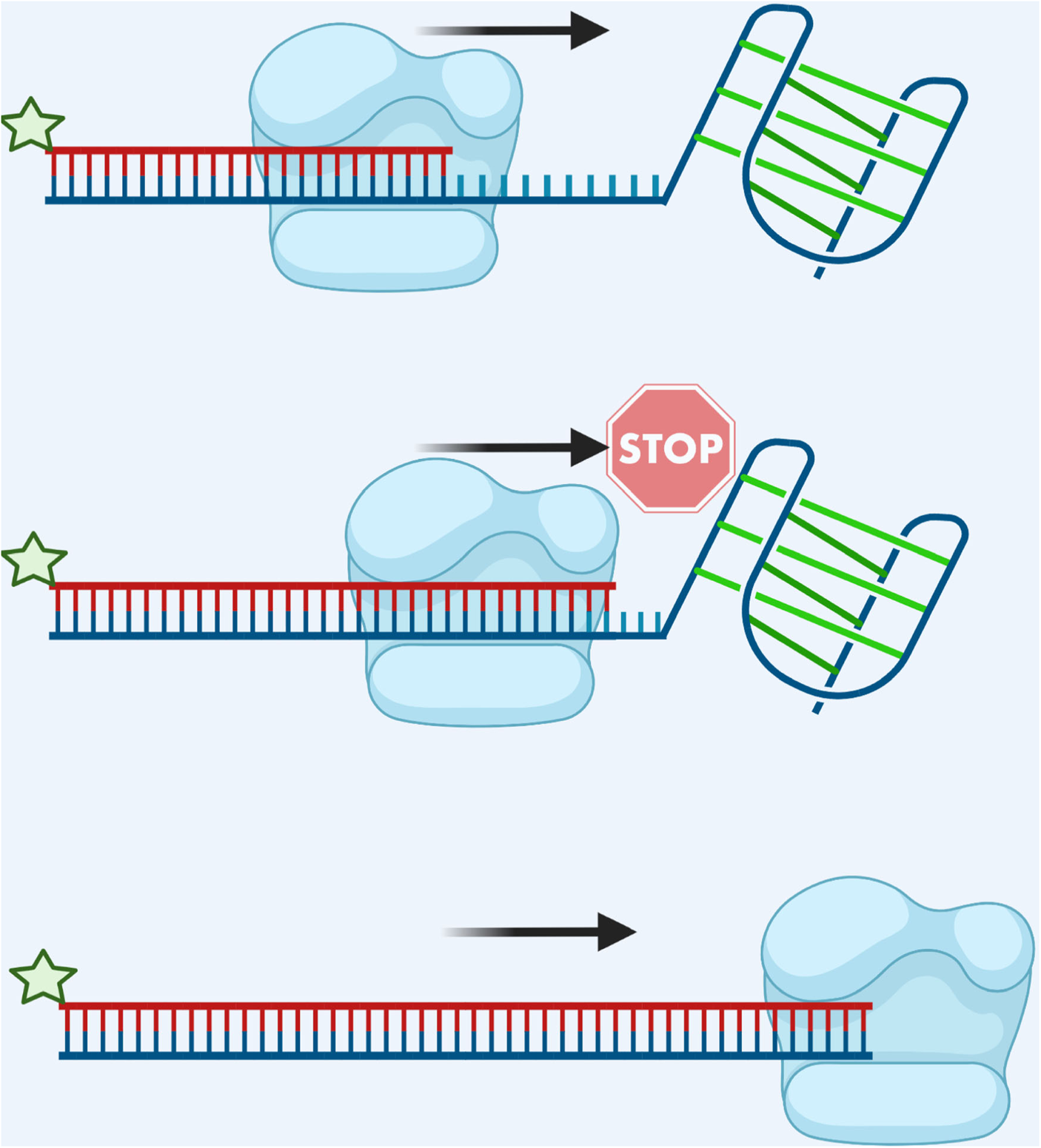
Scheme of high resolution primer extension assay. DNA polymerase binds template DNA containing folded iM (top). DNA primer is elongated until DNA polymerase meets iM, which blocks its progresion (middle). The DNA polymerase bypasses the unfolded iM (bottom). Created with BioRender.com

## Supplementary information

**Table S1.**
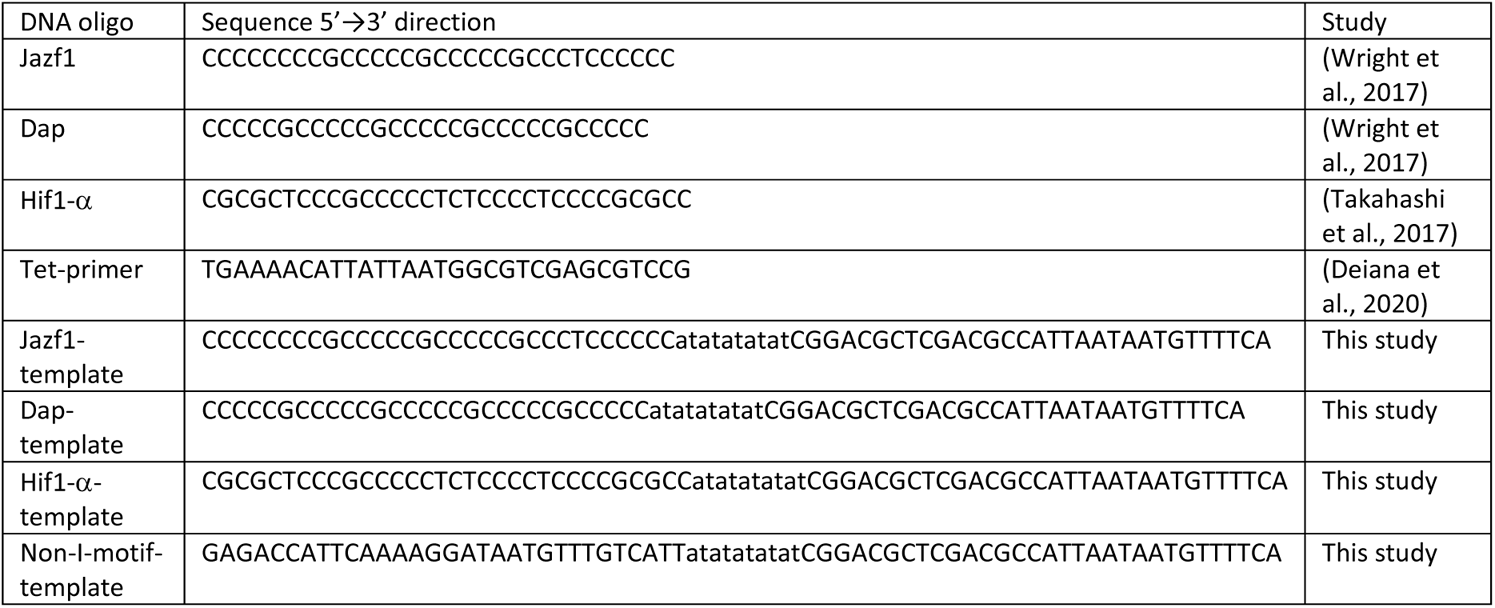
Oligonucleotides used in the study.

**Figure S1.**
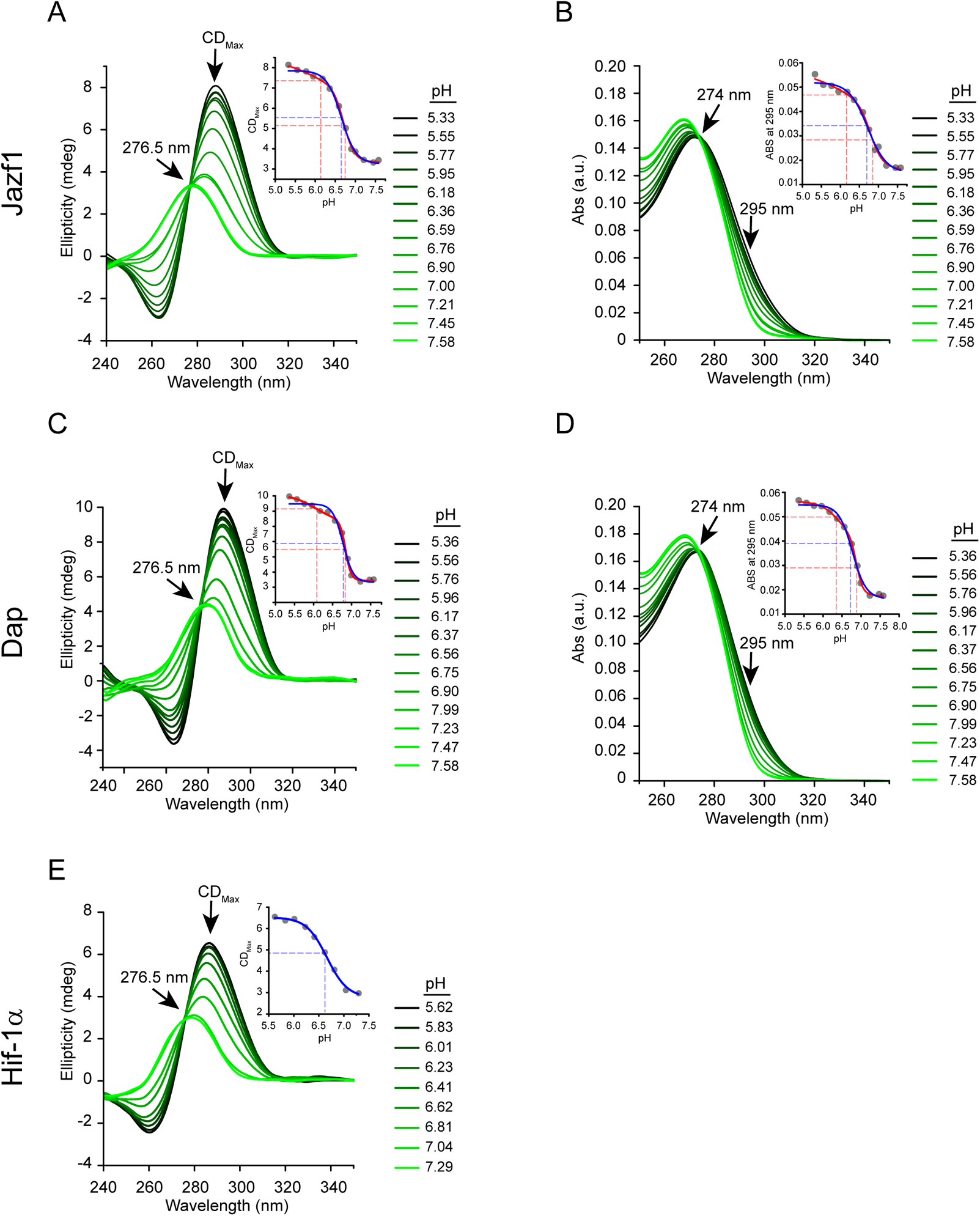
IMs fold at close-to-neutral pH at 25°C. CD and absorption spectra of iMs recorded at various pH: CD (A) and absorption (B) spectra of Jazf1, CD (C) and absorption (D) spectra of Dap, CD spectra of Hif-1**α** (E). Upper right corner of each graph contains CD_max_ or ABS_295_ values plotted over pH. These data were fitted into dose-response (blue) or bi-dose-response function (red) in OriginLab software for a visual comparison. Presented data are from one representative experiment. Final values are listed in Figure 1 and Figure S2.

**Figure S2.**
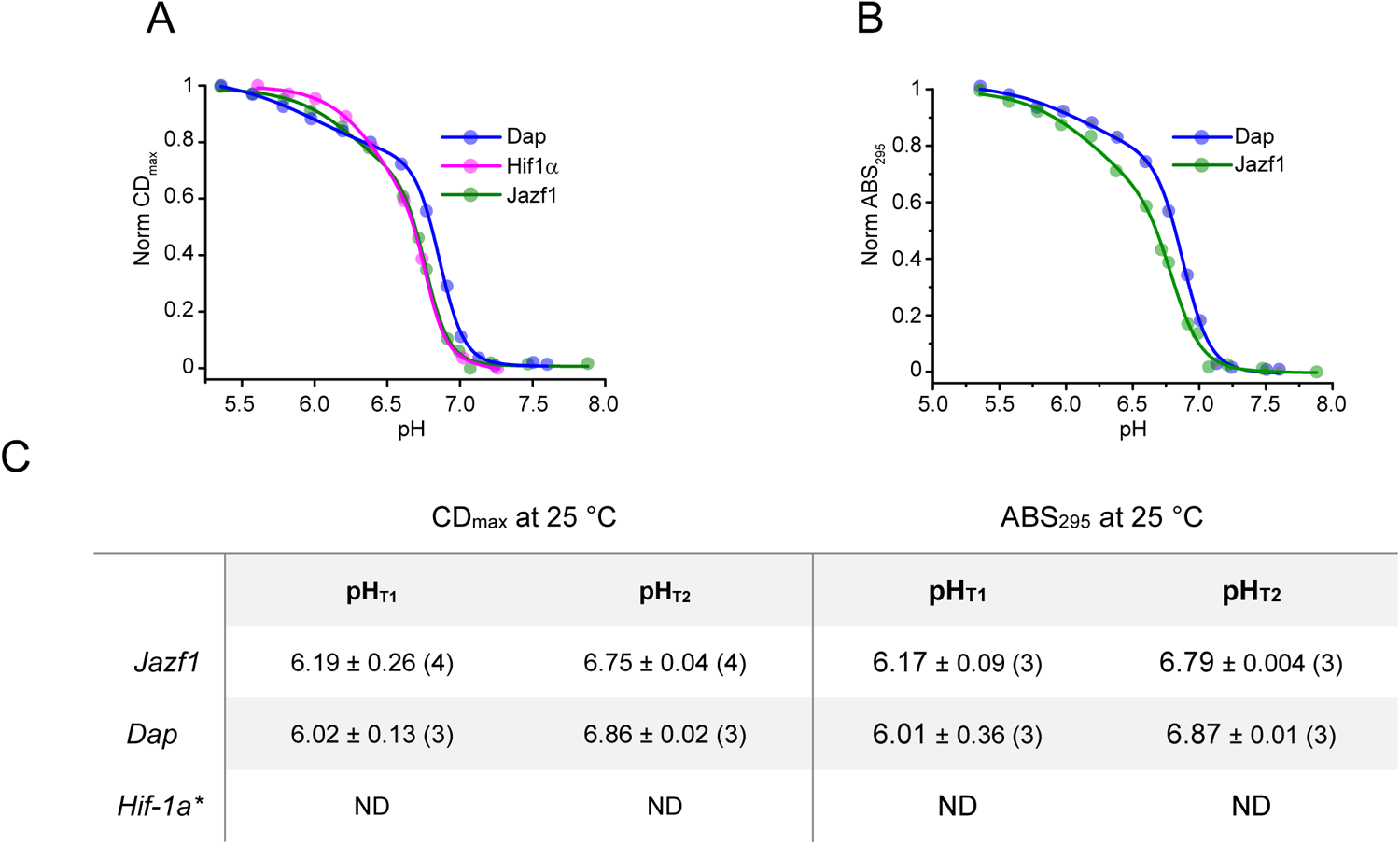
IMs fold at close-to-neutral pH at 25°C. CD_max_ (A) and ABS_295_ (B) of iMs spectra recorded at 25 °C. Data were fit into bi dose-response function to obtain two pH_T1_ and pH_T2_ values from two slopes. pH_T1_ and pH_T2_ values are listed in table in C. CD and absorption data were averaged and fitted in OriginLab software for visualization purpose only. * Fitting not successful/data missing.

**Figure S3.**
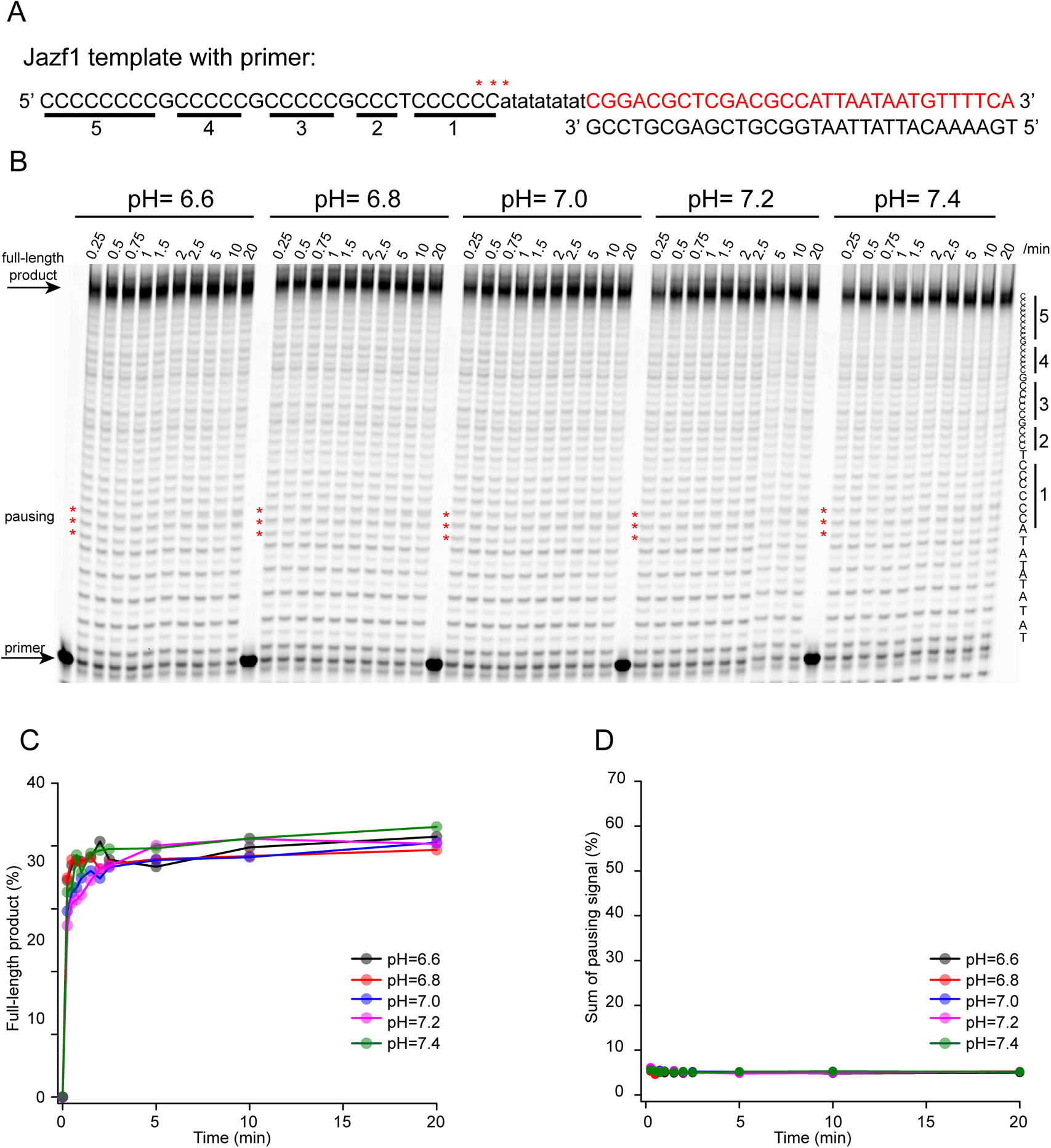
Jazf1 iM does not pause the DNA polymerase above pH=6.6 during DNA synthesis in primer extension assay. Sequence (A) and representative sequencing gel (B) of primer extension assay with template containing Jazf1 iM sequence, pH of the reaction is written above each gel. Time-points when individual samples were collected is written above each well. C-tracts from iM sequence are underlined in A and depicted on the right side of the gel. Pausing site is depicted as red stars (***) in each reaction and shown also in A. Quantification of full-length product (C) and pausing signal (D) of each reaction is describing an effect of pH on overall reaction and iM, respectively.

**Figure S4.**
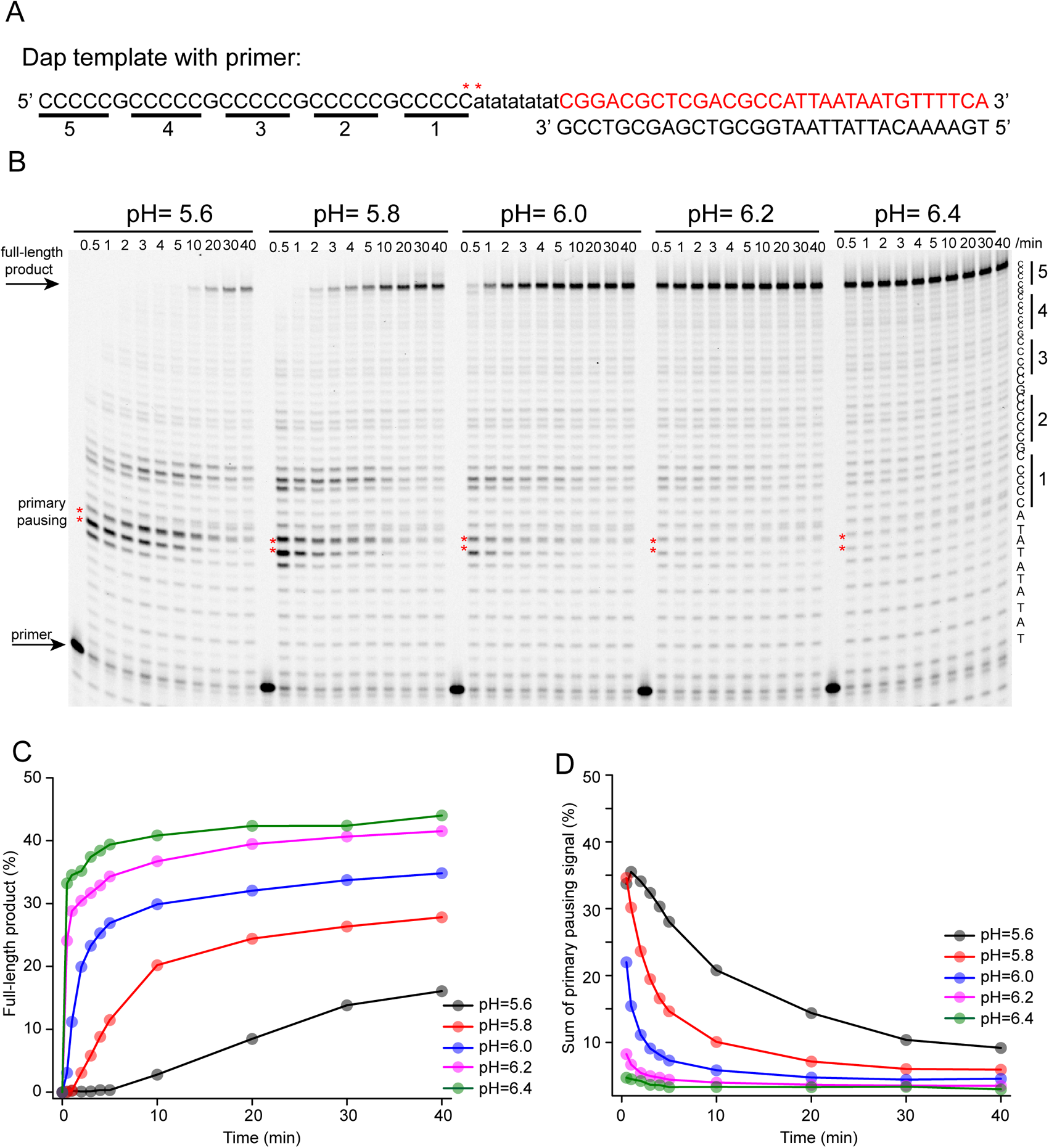
Dap iM pauses DNA synthesis in pH=5.6-6.0. Sequence (A) and representative sequencing gel (B) of primer extension assay with template containing Dap iM sequence, pH of the reaction is written above each gel. Time-points when individual samples were collected is written above each well. C-tracts from iM sequence are underlined in A and depicted on the right side of the gel. Main pausing sites are depicted as red stars (**) in each reaction and shown also in A. Quantification of full-length product (C) and the sum of pausing signal (D) of each reaction is describing an effect of pH on overall reaction and iM, respectively.

**Figure S5.**
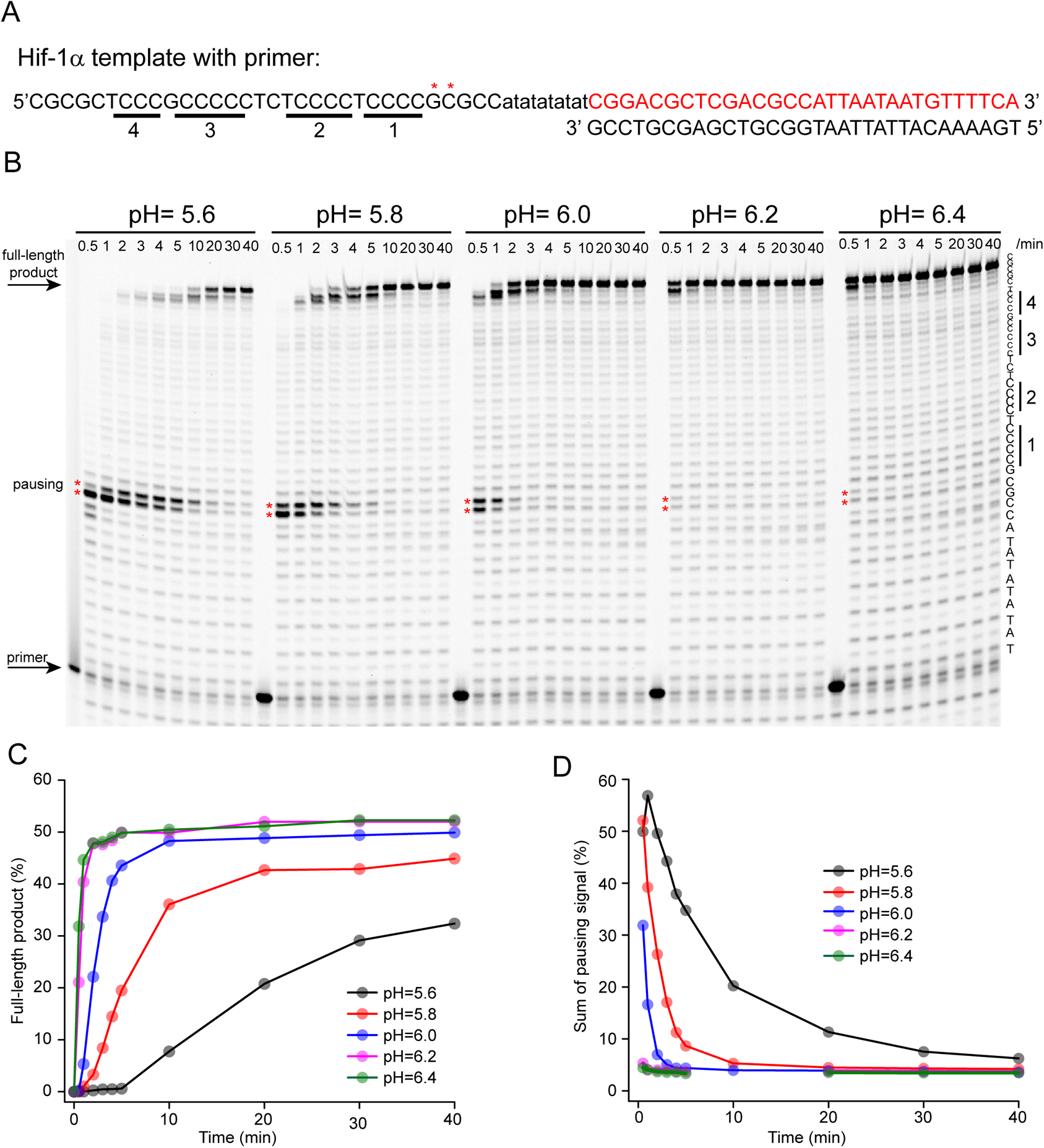
Hif-1α iM pauses DNA synthesis in pH=5.6-6.0. Sequence (A) and representative sequencing gel (B) of primer extension assay with template containing Hif-1**α** iM sequence. pH of the reaction is written above each gel. Time-points when individual samples were collected is written above each well. C-tracts from iM sequence are underlined in A and depicted on the right side of the gel. Pausing site is depicted as red stars (**) in each reaction and shown also in A. Quantification of full-length product (C) and the sum of pausing signal (D) of each reaction is describing an effect of pH on overall reaction and iM, respectively.

**Figure S6.**
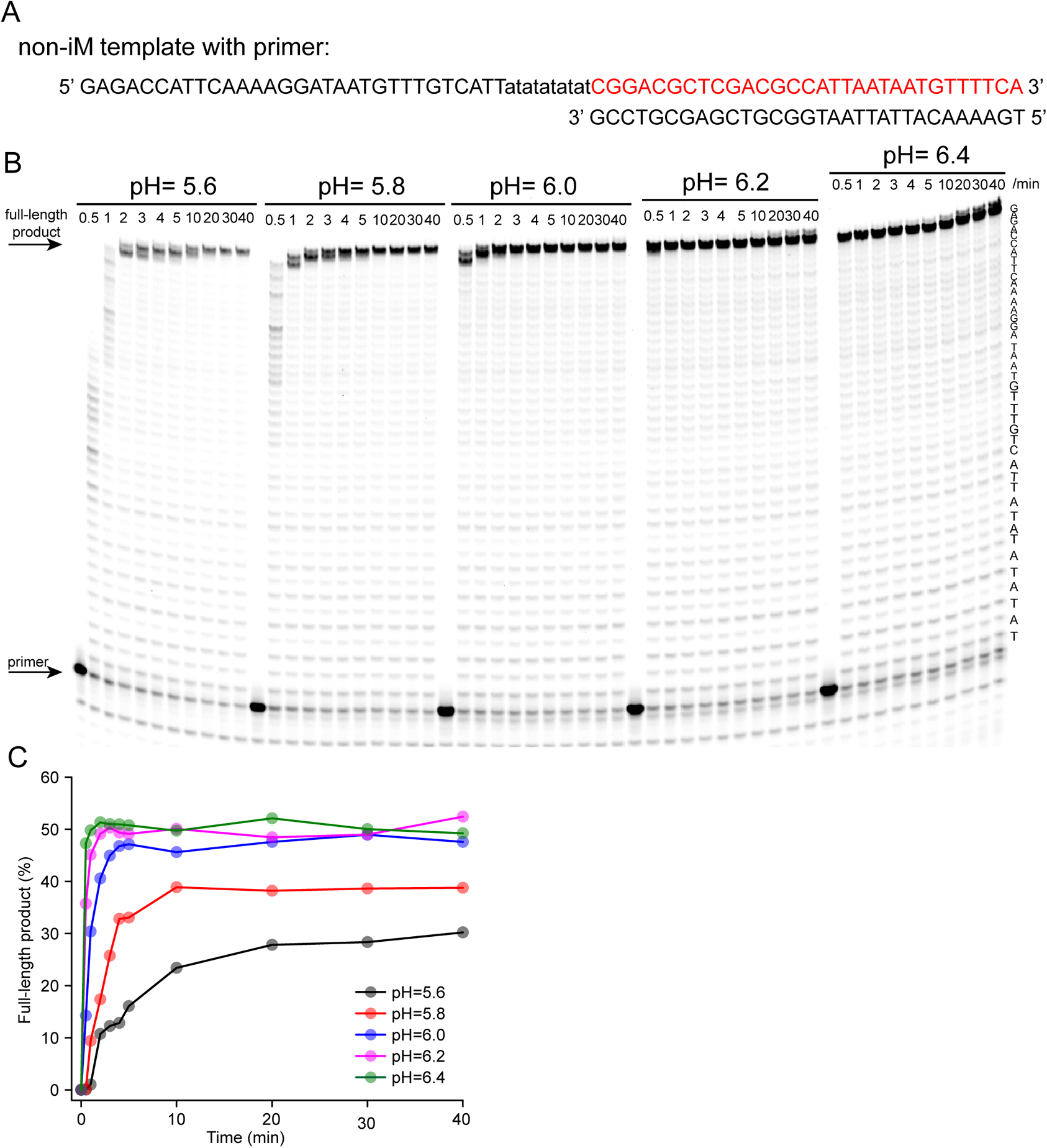
Non-iM control DNA with no secondary DNA structures allow DNA synthesis in pH=5.6-6.4. Sequence (A) and representative sequencing gel (B) of primer extension assay with ssDNA template (non-iM) sequence. pH of the reaction is written above each gel. Time-points when individual samples were collected is written above each well. Quantification of full-length product is describing an effect of pH on overall reaction.

**Figure S7.**
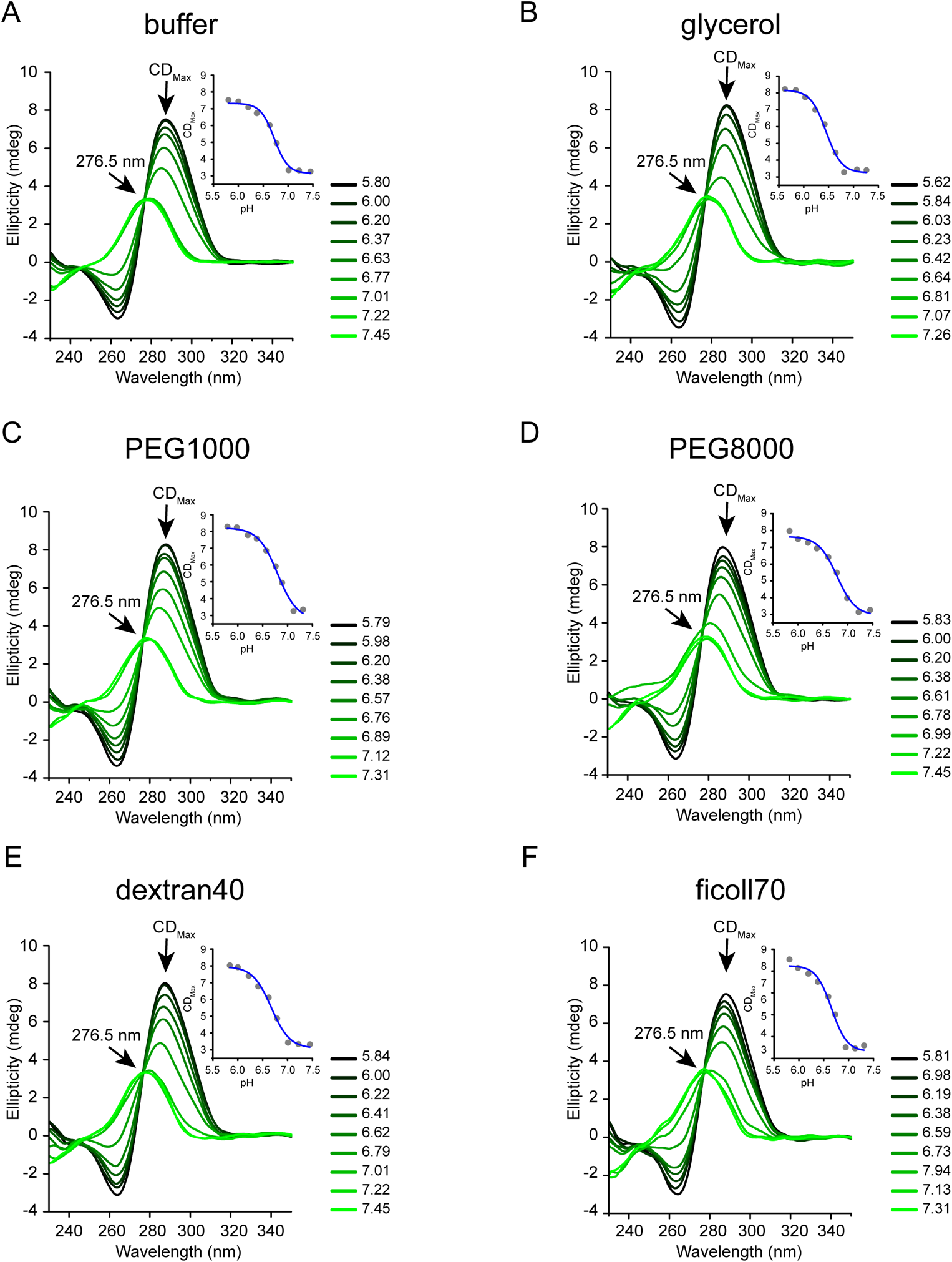
Molecular crowders and co-solutes do not shift pKa of Jazf1 iM into neutral pH. Individual CD spectra of Jazf1 iM in the presence of 20% co-solutes and molecular crowders: no additives (A), glycerol (B), PEG1000 (C), PEG8000 (D), dextran40 (E) and ficoll70 (F). CD spectra were recorded at 25 °C in different pH. CD_max_ values were plotted against pH and pH_T_ values were calculated by fitting the data into dose-response function (embedded graphs in each CD plot). pH_T_ values are graphically shown in Figure 1.

**Figure S8.**
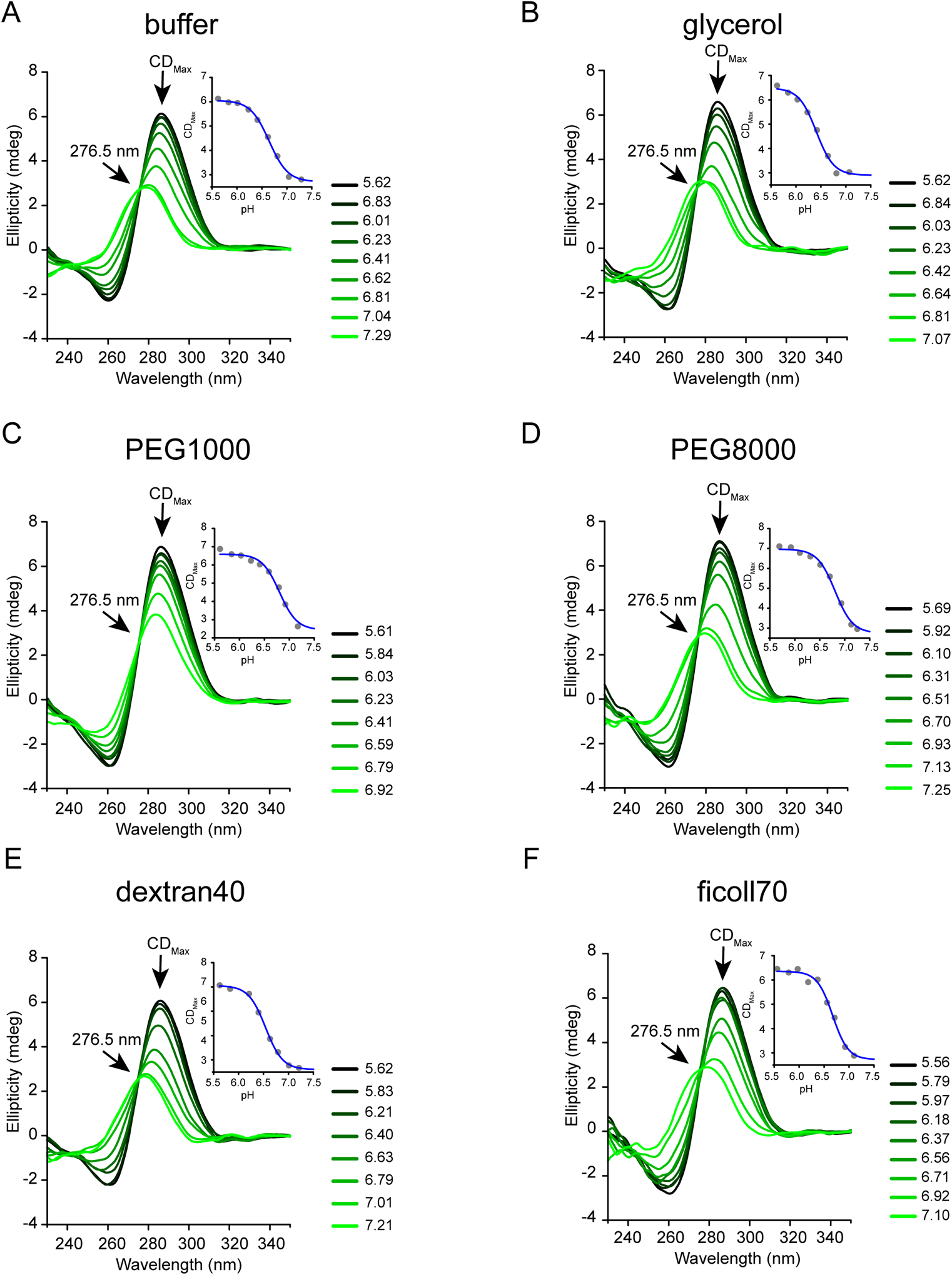
Molecular crowders and co-solutes do not shift pKa of Hif-1α iM into neutral pH. Individual CD spectra of Jazf1 iM in the presence of 20% co-solutes and molecular crowders: no additives (A), glycerol (B), PEG1000 (C), PEG8000 (D), dextran40 (E) and ficoll70 (F). CD spectra were recorded at 25 °C in different pH. CD_max_ values were plotted against pH and pH_T_ values were calculated by fitting the data into dose-response function (embedded graphs in each CD plot). pH_T_ values are graphically shown in Figure 1.

**Figure S9.**
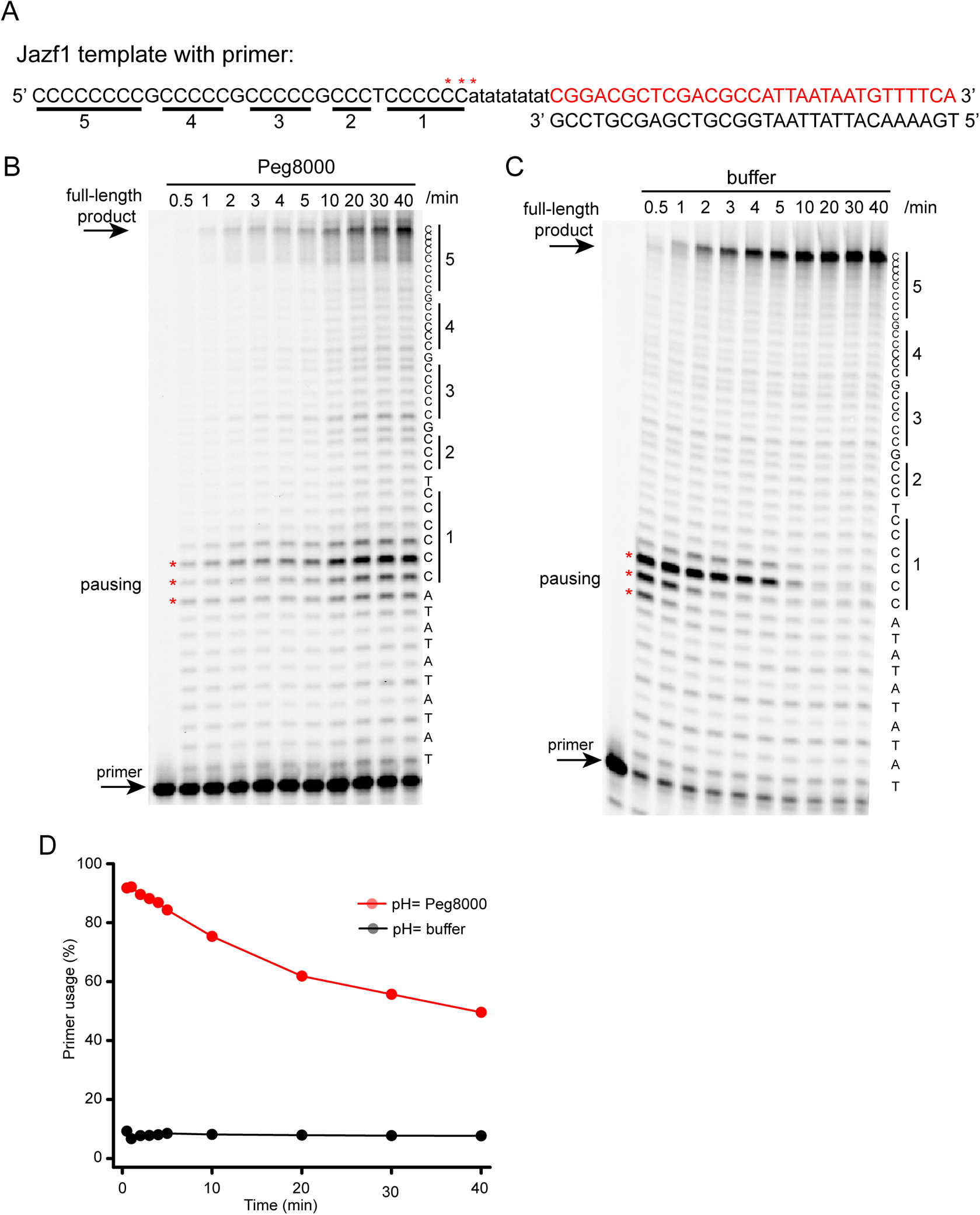
20% PEG8000 inhibits primer extension assay in an unspecific and non-iM dependent manner. Jazf1 sequence (A) and a sequencing gel (B) of primer extension assay supplemented with 20% PEG8000 and (C) with no supplements. Time-points when individual samples were collected is written above each well. C-tracts from iM sequence are underlined in A and depicted on the right side of the gel in B and C. Pausing site is depicted as red stars (***) in each reaction and shown also in A. Pausing site appearing in 10-40 minutes of reaction with PEG8000 cannot be a consequence of stabilized iM, but rather it is caused by very slow reaction initiation. D) Quantification of primer consumption from gels in B and C between 0.5-40 minutes.

**Figure S10.**
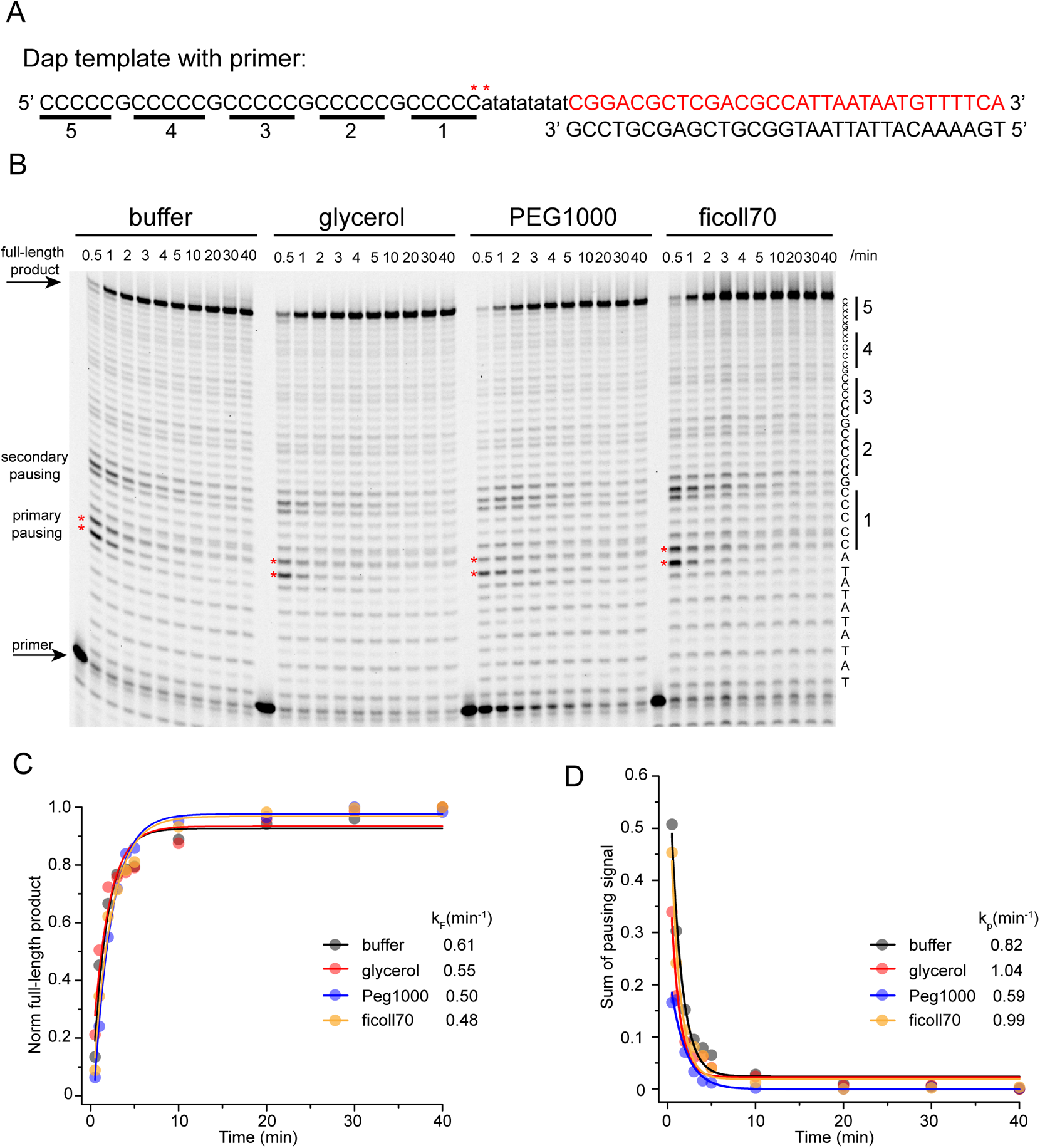
Molecular crowders and co-solutes do not stabilize Dap iM in primer extension assay in pH=6. Sequence (A) and representative sequencing gel (B) of primer extension assay with template containing Dap iM sequence run at 37 °C. Reactions were supplemented with 20% of co-solutes or molecular crowders. Specific condition is written above each gel. Time when individual samples were collected is written above each well. C-tracts from iM sequence are underlined in A and depicted on the right side of the gel. Pausing site is depicted as red stars (**) in each reaction and shown also in A. Quantification of full-length product (C) and pausing signal (D) of each reaction is describing an effect of additives on overall reaction and iM, respectively. Data from individual reactions are fitted into exponential equation in OriginLab software and apparent rate constants k_F_ and k_P_ are shown in C and D.

**Figure S11.**
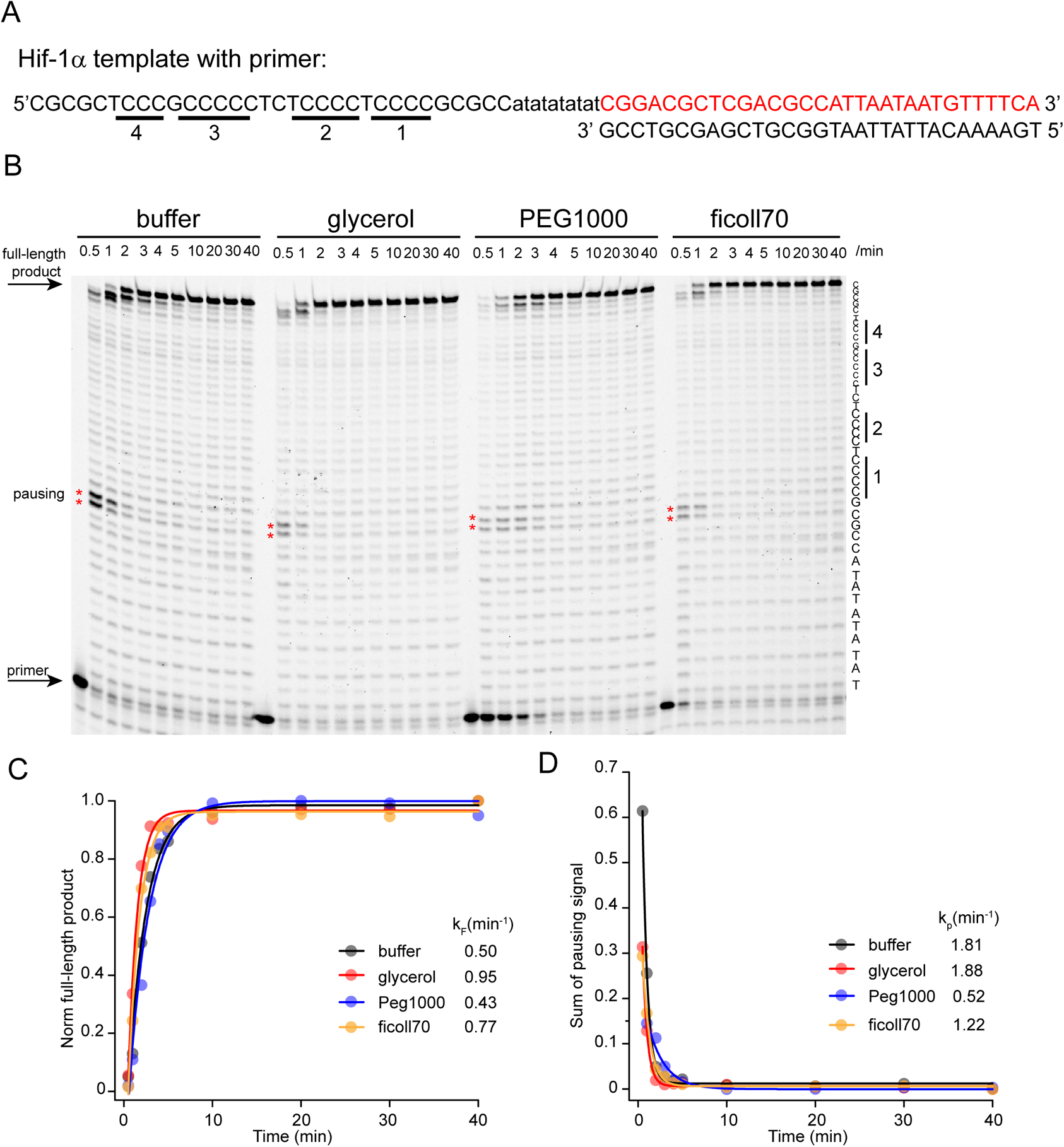
Molecular crowders and co-solutes do not stabilize Hif-1α iM in primer extension assay in pH=6. Sequence (A) and representative sequencing gel (B) of primer extension assay with template containing Hif-**α** iM sequence run at 37 °C. Reactions were supplemented with different co-solutes and molecular crowders. Specific condition is written above each gel. Time-points when individual samples were collected are written above each well. C-tracts from iM sequence are underlined in A and depicted on the right side of the gel. Pausing site is depicted as red stars (**) in each reaction and shown also in A. Quantification of full-length product (C) and pausing signal (D) of each reaction is describing an effect of additives on overall reaction and IM, respectively. Data from individual reactions are fitted into exponential equation in OriginLab software and apparent rate constants k_F_ and k_P_ are shown in C and D.

**Figure S12.**
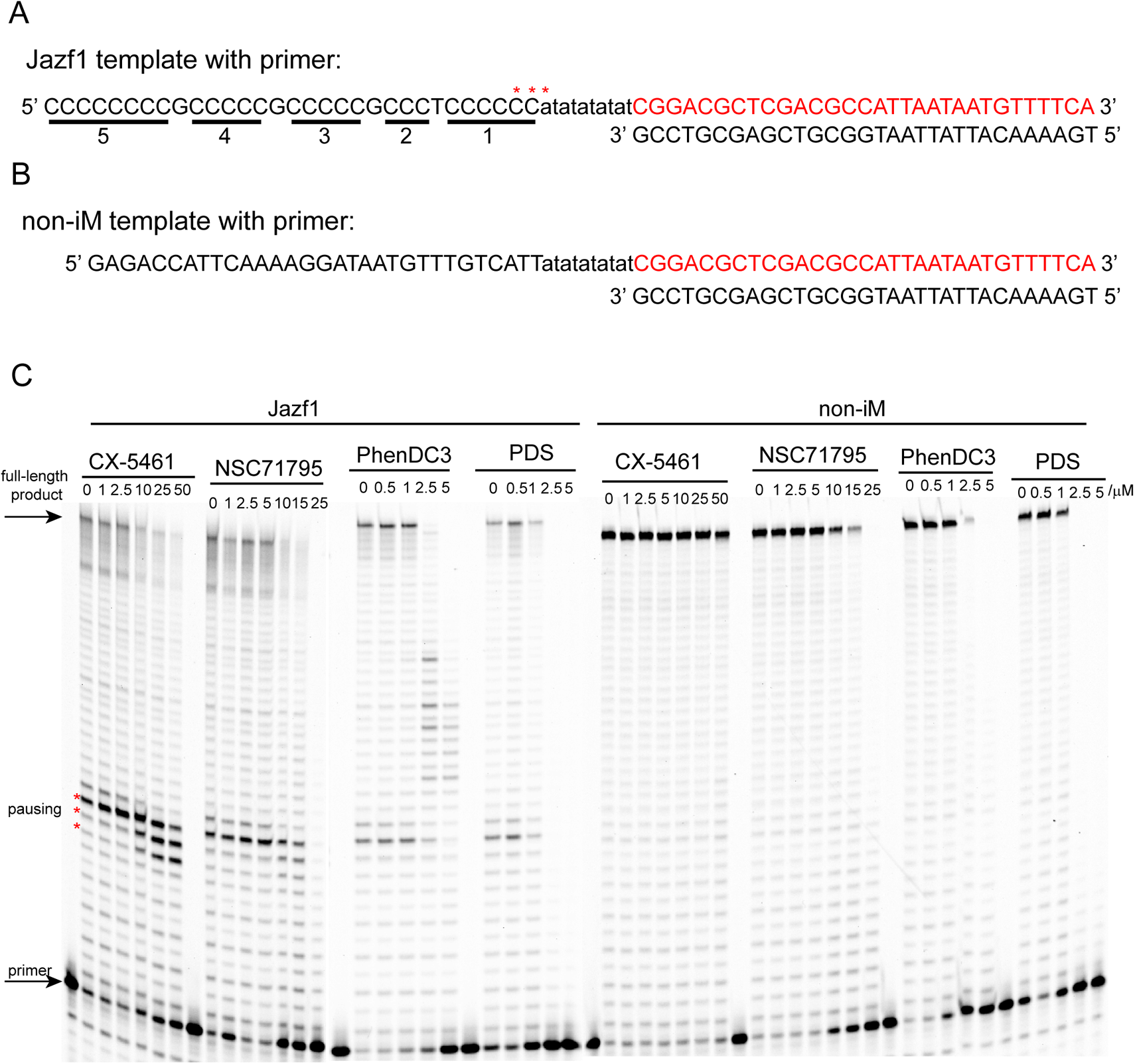
CX-5461 identified as a potential iM stabilizer in high resolution primer extension assay. Drug screening of sequences (A, B) and representative sequencing gel (C) of primer extension assay with templates containing Jazf1 iM and Non-iM sequence run for 4 min at 37 °C. Reactions were supplemented with various concentrations of small molecules (CX-5461, NSC71795, PhenDC3, Pyridostatin-PDS), described above each gel. C-tracts from iM sequence are underlined in A. Pausing site is depicted as red stars (***) in (A) and (C).

**Figure S13.**
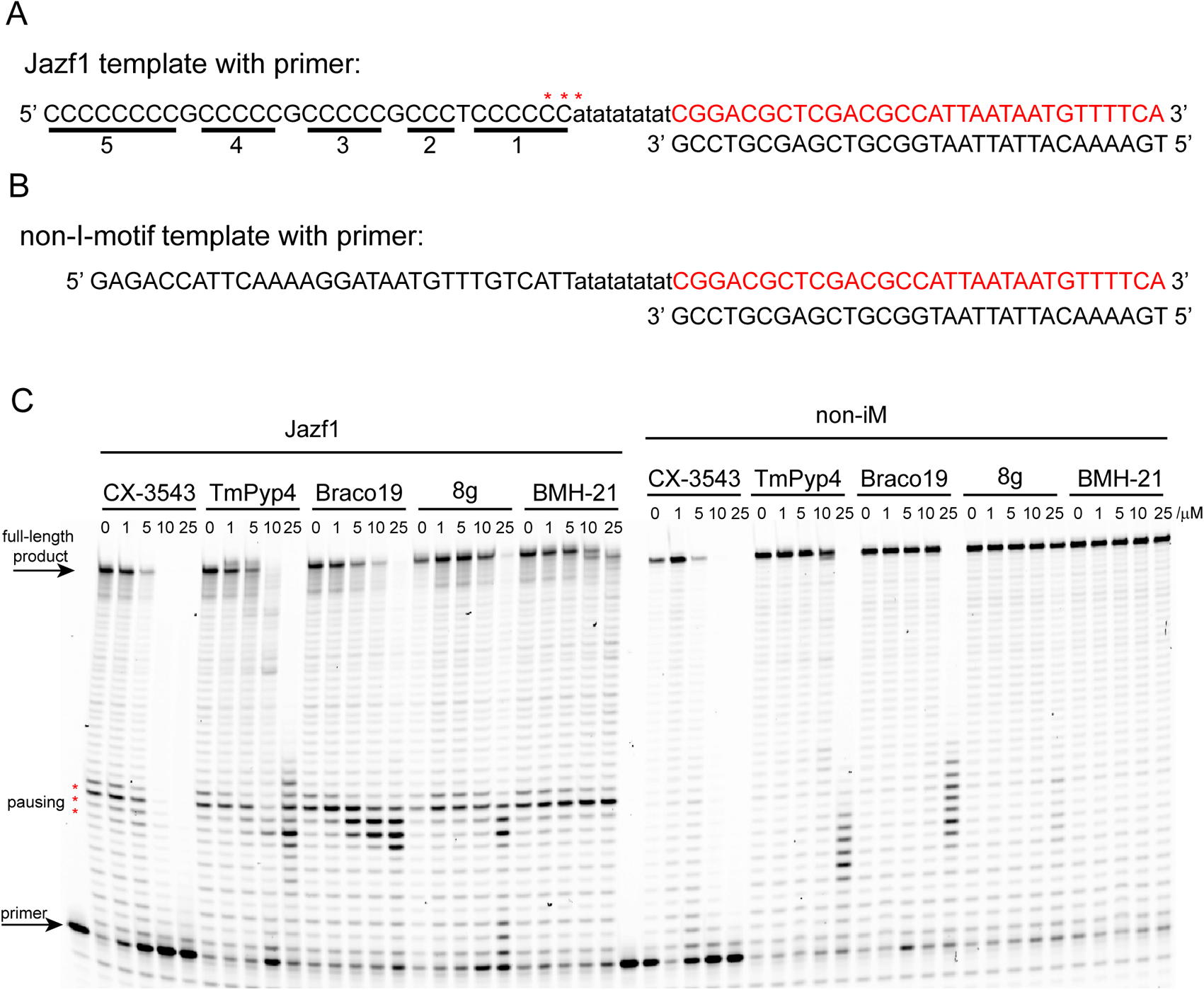
Ligands in high resolution primer extension assay. Sequences (A, B) and representative sequencing gel (C) of primer extension assay with templates containing Jazf1 iM and Non-iM sequence run for 4 min at 37 °C. Reactions were supplemented with various concentrations of small molecules (CX-3543, TmPyP4, Braco19, quinazoline 8g and BMH-21), described above each gel. C-tracts from iM sequence are underlined in A. Pausing site is depicted as red stars (***) in (A) and (C).

